# Unveiling Conserved Allosteric Hot Spots in Protein Domains from Sequences

**DOI:** 10.1101/2024.05.13.593877

**Authors:** Aysima Hacisuleyman, Dirk Fasshauer

## Abstract

The amino acid sequence determines the structure, function, and dynamics of a protein. In recent years, enormous progress has been made in translating sequence information into 3D structural information using artificial intelligence. However, because of the underlying methodology, it is an immense computational challenge to extract this information from the ever-increasing number of sequences. In the present study, we show that it is possible to create 2D contact maps from sequences, for which only a few exemplary structures are available on a laptop without the need for GPUs. This is achieved by using a pattern-matching approach. The resulting contact maps largely reflect the interactions in the 3D structures and contain information about its function and dynamics. This approach was used to explore the evolutionarily conserved allosteric mechanisms and identify the source– sink (driver-driven) relationships by using an established method that combines Schreiber’s concept of entropy transfer with a simple Gaussian network model. The validity of our method was tested on the DHFR, PDZ, SH3, and S100 domains, with our predictions consistently aligning with the experimental findings.

## Introduction

Proteins play a crucial role in cellular functions, often working in collaboration with other molecules to carry out tasks, and they are subjected to certain regulatory mechanisms, including allostery. Allostery refers to the regulation of a protein’s function by the binding of a regulatory molecule at a site other than the protein’s active site. The first allosterically regulated enzymes were discovered and characterized about half a century ago. It was discovered that allosteric binding can cause a change in conformation. The concept of a two-state conformational change was later extended, as reviewed by Kalodimos and Edelstein (Charalampos Kalodimos, 2013, July; Charalampos Kalodimos, 2013, May; Wu, Barahona and Yaliraki, 2024), so it is now assumed that allostery is best described as an ensemble of dynamic states.

Allosteric activation serves as a robust switch, toggling between an inactive and an active state. Typically, for allosteric mechanisms, multiple lower energy regions exist within a free energy landscape (Tsai and Nussinov, 2014). Mutations often disrupt this equilibrium, diverting the course of an allosteric communication pathway (Liu and Nussinov, 2016). Moreover, allosteric sites often serve as binding sites for drugs and are therefore of high pharmacological interest. The concept of “dynamic allostery,” as proposed by Cooper and Dryden, explains how allosteric communication pathways revolve around a mean conformational state, involving correlated thermal fluctuations and highlighting the entropic nature of allostery (Cooper and Dryden, 1984).

Dynamic allostery is a fundamental concept in the regulation of proteins’ function, where proteins undergo conformational changes at remote sites to respond to environmental cues. These remote interactions are essential for fine-tuning the proteins’ activity. Examples include hemoglobin’s ability to switch between states of binding and releasing oxygen (Ahmed, Ghatge and Safo, 2020), allosteric enzymes such as aspartate carbamoyltransferase adjusting their conformation depending on the availability of specific molecules (Helmstaedt, Krappmann and Braus, 2001), G protein-coupled receptors transmitting signals across cell membranes through long-range conformational changes (Li, et al., 2023), and cyclic nucleotide-gated channels in neurons undergoing conformational changes to transduce chemical signals into electrical signals (Hu, Zheng and Yang, 2023). These examples highlight the ever-changing and adaptive nature of proteins, showcasing how dynamic allostery is central to their biological functions by involving conformational fluctuations driven by thermal motion and entropic effects.

A tool used to explore conformational fluctuations in proteins arose from the concept of transfer entropy, which was introduced by Schreiber. This is a measure of the flow of information between two time series (Schreiber, 2000). It has been useful for identifying the causal inferences of correlated thermal fluctuations in proteins from the trajectories of molecular dynamics simulation (Barr, et al., 2011; Corrada, Morra and Colombo, 2013; Garcia Michel, et al., 2021; Hacisuleyman and Erman, 2017; Kamberaj and van der Vaart, 2009) or those produced by coarse-grained approaches such as elastic network models, specifically the Gaussian network model (GNM) (Acar, et al., 2020; Altintel, et al., 2022; Ersoy, et al., 2023; Garcia Michel, et al., 2021; Hacisuleyman and Erman, 2017; Hu, et al., 2020). It has also been applied in a broad range of fields to estimate the causal relationships in finance, neuroscience, social networks, and climate science. The GNM provides a simplified yet effective way to model the dynamics of protein structures (Bahar, Atilgan and Erman, 1997; Katebi, et al., 2015; Sen, et al., 2006; Yang, Song and Jernigan, 2007). It is the simplest elastic network model focusing on the inter-residue contact topology of the proteins and capturing the collective motions and fluctuations with significantly reduced demand for computer processing. We recently formulated a fast and efficient method for calculating the entropy transferred between pairs of residues by combining Schreiber’s concept of transfer entropy with the dynamic version of the GNM (dGNM) (Hacisuleyman and Erman, 2017) and detecting allosteric hot spots in proteins, which are conserved regions within a protein which are crucial for regulating their activities. It is important to note that all proteins may indeed exhibit allosteric communication to some extent, but the impact of specific residues on this communication can vary significantly(Ma and Nussinov, 2014; Reynolds, McLaughlin and Ranganathan, 2011).

Finding allosteric sites in a protein usually requires time-consuming biochemical, biophysical, and structural experiments. In recent decades, computer simulations of protein dynamics have played an increasingly important role. So far, only the dynamics of existing structures have been investigated using computer simulations. Such studies have been carried out selectively, and it has remained unclear to what extent an allosteric mechanism has been conserved in a given protein family. Compared with the ever growing number of protein sequences available in Uniprot (Consortium, 2023), the total number of high-resolution protein structures in Protein Data Bank (PDB) (consortium, 2019) is negligible. This bottleneck has made it difficult to gain insights into structural diversity. Methods of predicting structures have improved drastically over the years, shifting from the idea of identifying co-varying positions in multiple sequence alignments (MSAs) as directly interacting residues (Morcos, et al., 2011) to using deep neural networks. This bottleneck has widened substantially in the last few years, as programs are now available, such as Alphafold (Jumper, et al., 2021) and ESMFold (Lin, et al., 2023), with which the 3D structures of proteins can be predicted. To predict the structures, these programs make use of physical, geometric, and evolutionary constraints. The evolutionary constraints used are based on the fact that homologous sequences fold into similar structures. MSAs show that protein families frequently possess extended segments of conserved residues, which play a critical role in the protein’s function. For instance, cell surface receptor proteins exhibit diverse amino acid sequences at their binding sites for extracellular cues, yet they display greater similarity in their amino acid sequences that engage with common intracellular signaling proteins.

To comparatively investigate the dynamics of entire protein families, a variety of predicted structures can now be included in analyses. However, examining an entire protein family could prove overly resource-intensive in terms of both the computational and memory requirements, as each structure would have to be calculated first. A more straightforward and computationally efficient approach for deriving structural attributes from sequences involves generating their contact maps. A contact map serves as a 2D depiction of the relative closeness of residues within the protein’s spatial arrangement. This, in turn, can be used to investigate the structural dynamics of the protein using GNMs. Tirion (Tirion, 1996) demonstrated that a single parameter harmonic potential could effectively capture the large amplitude motions of proteins in their native states through normal mode analysis. Building upon these principles Haliloglu, Bahar and Erman(Haliloglu, Bahar and Erman, 1997) proposed a model, which can adopt a single parameter for the harmonic potential and predict the equilibrium fluctuations of the alpha carbons. This model requires the connectivity matrix, as known as the Kirchhoff matrix, which can be derived from the contact maps. By performing singular value decomposition and computing the inverse of the Kirchhoff matrix, one can find the square fluctuations, which are the diagonal elements of the inverse Kirchhoff matrix. In addition, the cross-correlations between atomic fluctuations can be calculated by using the eigenvectors and eigenvalues of the Kirchhoff matrix, providing a detailed view of the collective motions within a protein. Individual modes from the collective motions can be studied separately and further classified into slow and fast modes of motion. Slow modes of motion represent collective motions, often involving coordinated movements of multiple structural elements and are typically associated with biologically relevant processes, such as conformational changes and hinge movements. Whereas fast modes of motion represent high-frequency vibrations that usually correspond to smaller, more localized movements within the molecule, such as side-chain vibrations. Slow and fast modes are associated with function and stability, respectively (Bahar, et al., 1998; Bahar, et al., 2010; Jernigan, Demirel and Bahar, 1999).

As mentioned, MSAs, or individual protein sequences, also contain information on the residues that are in direct contact. This information can therefore also be used to create contact maps without having to calculate the complete 3D structure of the protein. It is, in principle, possible to go directly from the sequences to the functional dynamics of a protein using direct coupling analysis (DCA), for example, as has recently been shown (Jia, Kilinc and Jernigan, 2023). Such shortcuts mean an immense saving in computational power, given the huge volume of protein sequences available.

In this study, we have taken a different approach to tap into the vast amount of sequence information available to generate contact maps, providing a simple yet effective means to investigate the complex landscape of proteins’ functions and allosteric regulation. We developed a pattern-matching approach, which identifies short, conserved sequence motifs from homologous 3D structures from PDB (consortium, 2019) or AlphafoldDB (Schaeffer and Daggett, 2011; Varadi, et al., 2022). These sequence motifs are used to generate contact maps for given protein sequences. The contact maps are then used to apply the entropy transfer concept of Schreiber(Schreiber, 2000) in combination with dGNM (Hacisuleyman and Erman, 2017) to identify sources and sinks of information, as previously described. As an addition, the square fluctuations and the cross-correlations are calculated from the contact map derived Kirchhoff matrices and are compared with the experimental structures. The cross-correlations at the slow modes of motions are used to support the detected source and sink residues in terms of their importance for the functionality. They provide crucial information about the coupled motions of different parts of a protein. While square fluctuations tell us about the overall mobility of individual residues, cross-correlations show how these motions are interconnected. This information is crucial for understanding the function, allosteric communication and stability.

This methodology offers a computationally efficient means to pinpoint potential allosteric regulation sites, opening new avenues for understanding the conserved allosteric regulation mechanisms in protein domains and relating them to the proteins’ functions and study the dynamics of the system in a fast and efficient way. We considered four protein domains, namely DHFR, PDZ, S100, and SH3, which are widely studied for their allosteric behavior, and determined a common sink–source behavior for each domain.

## Methods

### Collecting sequences

First, we extracted the sequence datasets for the following protein domains from the Pfam database (Mistry, et al., 2021) using the representative Proteome 75 (Chen, et al., 2011) dataset: dihydrofolate reductase (DHFR) (Pfam ID: PF00186), PSD-95/Discs-large/ZO-1 (PDZ) (Pfam ID: PF00595), SRC Homology 3 (SH3) (Pfam ID: PF100018), and S100 (Pfam ID:

PF01023). We selected these four widely studied protein domains in order to establish our method. For each of the domains, high-resolution structures are available. From the initial set of sequences from Pfam (Mistry, et al., 2021), we removed any duplicate sequences to prevent redundancies. For each sequence, the conserved domain was extracted using the appropriate Hidden Markov model (HMM) (Johnson, Eddy and Portugaly, 2010) specific for that domain downloaded from Pfam(Mistry, et al., 2021). We performed an HMM search and considered the sequences with an alignment score higher than the overall mean bit-score and a length greater than or equal to the respective HMM motif, which covered the entire sequence region for the specific domain. Initially, we downloaded 43,000 DHFR, 173,969 PDZ, 93,857 SH3, and 5,082 S100 domain sequences. After the selective filtering step, we established a set of high-quality sequences. The number of sequences per domain was 5,895 for DHFR, 5,070 for PDZ, 4,674 for SH3, and 2,507 for the S100 domain.

### Generation of contact maps: pattern search and matching

For generating contact maps for the set of sequences, we searched the PDB (consortium, 2019) using the JackHMMER (HMMER 3.4) (Johnson, Eddy and Portugaly, 2010) tool for the structures of each of the four studied domains. Note that for sequences that do not have homologous structures, AlphaFoldDB (Varadi, et al., 2022) could be utilized as an alternative resource. For each of the four protein families, we curated a list of homologous PDB structures. In these structures, we subsequently identified contact patterns, namely the center of mass of residues of up to four amino acids that were located at a maximum distance of 8 Å from each other. Conserved contact patterns were successful in identifying novel motifs in a specific set of protein domains in an earlier study (Bradley, Kim and Berger, 2002), as they represented a specific fold. All sequences in a set were then searched for homologous structures, and the contact patterns were extracted and saved for each domain. For each pattern, residues that were located 8 Å from each other were denoted in uppercase letters and the residues in between are denoted in lowercase letters. To generate the contact maps, we used MATLAB’s (MATLAB (R2023b)) *localalign* (Barton, 1993) function and locally aligned the uppercase elements within a pattern to the query sequence with a gap-opening penalty of 10, which was set to 8 by default. An empty N × N contact matrix, C, was generated where N is the length of the sequence. If three uppercase strings within the pattern aligned with the residues i, j, and k of the query sequence, a value of 1 was added to each corresponding pair within the contact matrix, namely, C(i, j), C(i, k), and C(j, k).

For each domain, the contact maps established by the pattern-matching approach were compared with contact maps generated directly from the established structures.

### dGNM and calculation of transfer entropy

GNM is a simplified representation of proteins’ dynamics and structural fluctuations. It is widely used to study the collective motions and essential dynamics of proteins. GNM provides insights into the low-frequency global motions of proteins by considering only the connectivity between atoms, without explicitly simulating the atomic interactions. The connectivity between the atoms is represented by the Kirchhoff matrix, Γ. The contact map can be used to define the Kirchhoff matrix. The Γ matrix and the contact map (CM) have the following format:

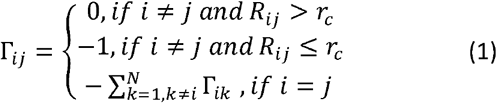

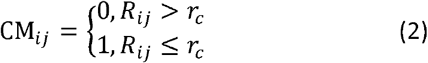

where i, j, and k represent the residue indices, R_ij_ is the distance between the Cα atoms, and r_c_ is the cutoff distance, taken as 8 Å for this study. The Γ matrix relates the correlations of residues to their connectivity.

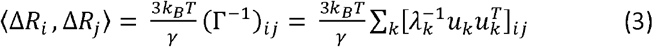

Where k_B_ is the Boltzmann constant, T is the temperature, the γ is the force constant, considered to be uniform for all, λ are the eigenvalues and u_k_ are the k^th^ eigenvectors. Since the Γ matrix is singular, we compute the inverse of it using singular value decomposition (SVD). This allows us to extract its eigenvalues and eigenvectors for the inversion process.

Schreiber’s transfer entropy formulation helps us understand how the past behavior of one process influences the future behavior of another process; therefore, it requires a time delay, τ, and describes the causal relationships between processes. For this study we used τ = 5 ps, since the characteristic time for the correlations to decay to their 1/e values is peaked around 5 ps. The dynamic version of the GNM describes the entropy transferred from residue i to residue j with the following equation:

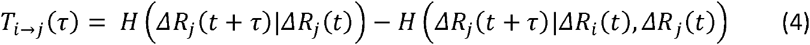

where the first term, *H* (*ΔR*_*j*_(*t* + τ)| *ΔR*_*j*_(*t*)), is the conditional entropy of residue j at time t+τ, given its values at time t. The second term, *H* (*ΔR*_*j*_(*t* + τ) | *ΔR*_*j*_(*t*), *ΔR*_*j*_(*t*)),, is the conditional entropy of residue j at time t+τ, given the values of residue i and j at time t. The result T_i→j_ (τ) indicates the extent of the reduction in the entropy in the future fluctuations of residue j, given the previous values of residues i and j. When the conditional entropy terms are expressed in terms of probabilities, commonly known as Shannon’s entropy formula, the transfer entropy equation becomes:

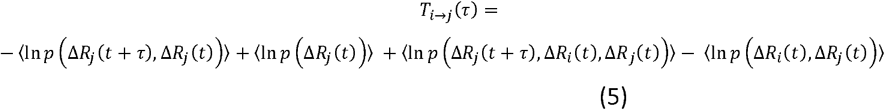

The time correlations of the residues’ fluctuations are given by the dGNM as:

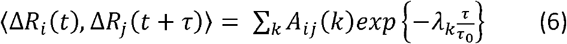

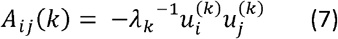

where *λ*_*k*_ is the k^th^ eigenvalue and 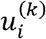 is the i^th^ component of the k^th^ eigenvector of the Γ matrix. The time τ_0_ is the characteristic of the vibrational dynamics of all proteins, and we and used τ_0_ = 1 ps. When Eqn. 6 is substituted into Eqn. 5, the transfer entropy formula becomes:

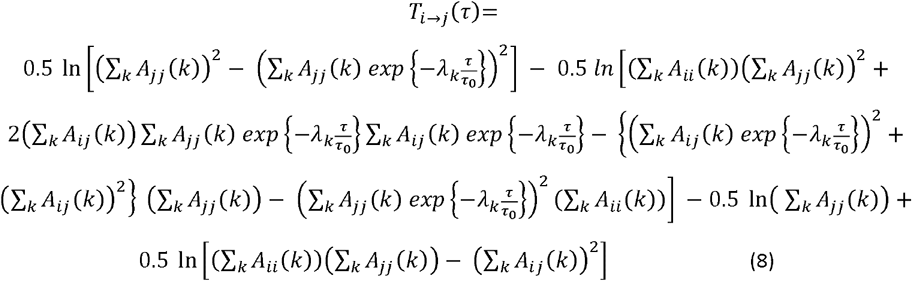

The details of the formulation can be found in previous articles (Erkip and Erman, 2004; Hacisuleyman and Erman, 2017; Haliloglu, Bahar and Erman, 1997) and the full derivation can be found in the supplementary material of (Hacisuleyman and Erman, 2017). We can use the net entropy transfer that accounts for a causal relationship to understand which residue is the driver and which residue is the driven. The net entropy transfer is the difference between the entropy transferred from residue i to j and the entropy transferred from residue j to i. We define net transfer entropy as:

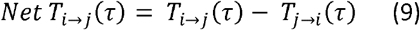

A positive net transfer entropy value indicates that residue i exerts an influence on the fluctuations of j, and vice versa. The net entropy transfer equals zero under two conditions: (1) when residue i and residue j are not correlated or (2) the transfer from residue i to j is the same as that from residue j to residue i, indicating no causality. When we combine the net entropy transfer from residue i to all available residues j, we can determine the overall behavior of residue i, particularly whether it acts as an entropy source or an entropy sink.

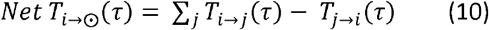

If the cumulative net entropy of residue i has a positive value, residue i is the source of entropy for the system, where information flows from residue i to the rest of the system. Conversely, if the cumulative net entropy of residue i has a negative value, information flows from the source residues to residue i, designating it as an entropy sink.

We performed a dGNM analysis on the predicted contact maps derived from protein sequences. Subsequently, we calculated the net transfer entropy for each protein and used Eqn. 10 to plot the cumulative net entropy transfer values and identified local maxima on the positive side and local minima on the negative side of the plot.

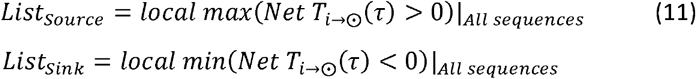

We stored the cumulative net entropy transfer values for these local maxima in a list of “sources” and those for the local minima in a list of “sinks” with their cumulative net entropy values.

After generating the “source” and “sink” lists for a protein family, we summed the collected cumulative net entropy transfer values over all sequences. By summing these values across all sequences, we gained insights into each residue’s role in the allosteric communication within the domain. This method helped us find conserved entropy source and sink regions in the protein domains. To be able to identify the exact source and sink residues for protein families, we set thresholds for positive and negative values. We used a quantile-based approach. The threshold for the sources was set at the 85^th^ percentile, and the threshold for the sinks was set at the 15^th^ percentile.

### Square Fluctuation and Cross-correlation Analysis by using GNM

To analyze the square fluctuations of our protein sequences, we used GNM. We generated the Γ matrix by repurposing the predicted contact maps for all sequences as shown in Eqn.

1. Each diagonal element of the Γ matrix characterizes the local packing density of residues. The inverse of the Γ can be written as

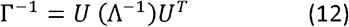

Where the columns of U matrix are the eigenvectors and Λ is a diagonal matrix of the eigenvalues. Mean square fluctuations of the Cα atoms and the cross-correlations between their fluctuations are determined from the diagonal off-diagonal elements of the Γ^-1^ matrix, respectively, as shown in Eqn. 3. By using the normal mode, square fluctuation(*calcSqFlucts*) and cross-correlation calculation(*calcCrossCorr*) functions of the ProDy package (Bahar, Atilgan and Erman, 1997; Bakan, Meireles and Bahar, 2011; Haliloglu, Bahar and Erman, 1997) we calculated the square fluctuations from our generated Γ matrices, for the first 10 normal modes, and we reported the mean square fluctuations for our sequences. As for the PDB structures, we extracted the B-factors from their PDB files and plotted the mean values. Since the in Eqn. 3 is not set to scale with the experimental temperature factors and taken as 3*k*_*B*_*T*, the GNM fluctuations are smaller than the temperature factors (Haliloglu, Bahar and Erman, 1997).

For the cross-correlations, we generated cross-correlation plots for the contact map derived Kirchhoff matrices for the first 1, 3 and 10 modes. The plots were calculated for each sequence and then we computed the mean cross-correlations by averaging over all cross-correlation matrices, and we reported the mean cross-correlation plots. Similarly, for the experimental structures, we calculated the cross-correlation plots for modes 1, 3 and 10 and reported the mean plots. To quantify the difference between the mean generated cross correlations with the mean cross correlations of the experimental structures, we used the mean-squared error (MSE).

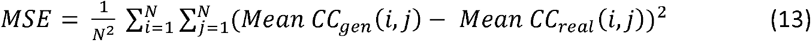

Where N is the number of residues, mean CC_gen_ is the mean cross correlations of the contact map derived Kirchhoff matrices and mean CC_real_ is the mean cross correlations of the available structures.

## Results and discussion

Proteins are classified into protein families on the basis of their sequences and structural similarities, which have a common evolutionary origin. This can be used to study their function and allosteric regulation. We have developed a method to quickly convert the sequences of protein families into contact maps without having to model a 3D structure for each sequence. To test our approach, we selected domains with different folds. We used four protein domains for which high-resolution structures are available and whose allosteric behavior has been sufficiently studied, namely the DHFR, PDZ, SH3, and S100 domains. This allowed us to use high-quality sequences from the Pfam database. To use the available high-resolution 3D structures for the domains, we recursively searched the PDB using JackHMMER. The characteristic folds of these protein domains are often established through characteristic conserved sequence motifs that are in direct contact in the 3D structure. We applied a contact threshold of 8 Å and selected characteristic patterns of up to four amino acids. Contact patterns have been used successfully in identifying novel motifs in a specific set of protein domains (Amala and Emerson, 2019; Bradley, Kim and Berger, 2002). We have extended this strategy by using the contact patterns found this way to establish the contact map. The contact maps were then transformed into the Kirchhoff matrix and used in the dGNM for further analysis. For each protein, we compared a contact map established from the available PDB structures with the contact map established solely with our pattern-matching approach. In all cases, the distant interactions in the 3D structure were clearly recognized by the pattern-matching approach, which allowed us to study the protein domain in detail without having the high-resolution structures available. Our approach is illustrated in Fig. 1.

**Figure 1.**
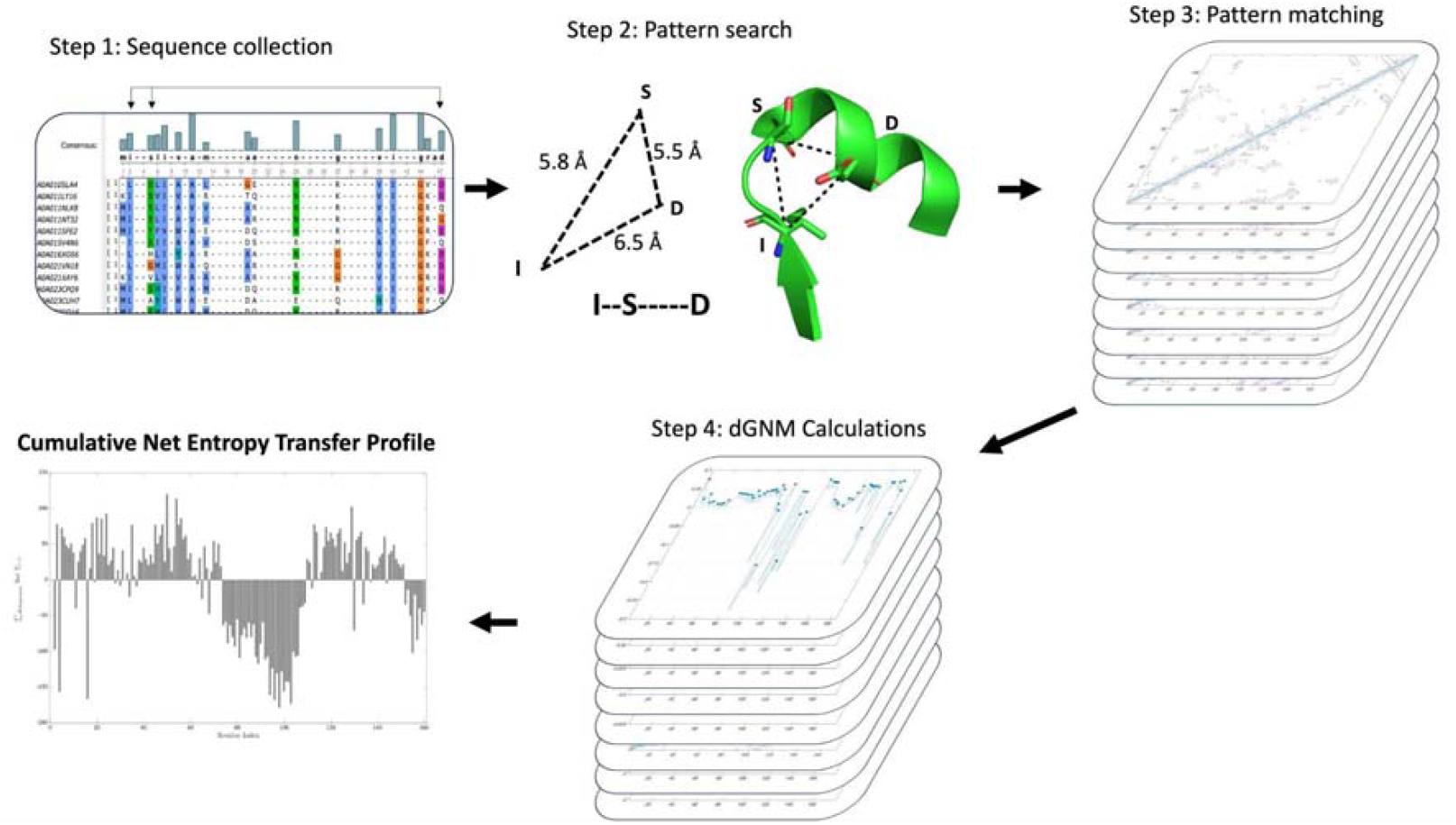
Workflow illustrating the procedure for finding the net entropy transfer profiles for protein domains without their high-resolution structures being available. In the first step, the sequences of a protein family were searched for with already established HMMs in the Pfam database, and a non-redundant set was generated. In Step 2, contact patterns were established with the help of homologous protein structures. In Step 3, these contact patterns were searched for in the individual sequences, and a contact map was created. These contact maps were then used for dGNM calculations and finding the net entropy transfer profile for specific protein domains (Step 4).

### DHFR

The enzyme DHFR converts 7,8-dihydrofolate (DHF) into 5,6,7,8-tetrahydrofolate, an important cofactor in many enzymatic reactions, by using nicotinamide adenine dinucleotide phosphate (NADP) as a hydride donor. Its allosteric mechanism has been studied extensively via a variety of experimental and computational approaches (Goldstein and Goodey, 2021). The backbone of the DHFR domain, which is approximately 160 residues long, consists of a central beta-pleated sheet composed of eight strands. Of these strands, seven align in a parallel, whereas the eighth adopts an antiparallel orientation. These beta strands are joined together by four alpha helices. Three loops play a significant role in the function: the Met20 loop (Residues 9–23), the βF-βG loop (FG loop; Residues 116–132), and the βG-βH loop (GH loop; Residues 135–149). The active site is located within the Met20 loop, which features a conserved Pro-Trp dipeptide at Positions 21 and 22, respectively. The tryptophan within this dipeptide plays a crucial role in the substrate-binding of the enzyme. DHFR switches its conformation during its catalytic cycle, going between an open and a closed state, depending on whether its active site is available or blocked by the Met20 loop. The residues that are involved in the allosteric communication pathway have been identified by nuclear magnetic resonance experiments, molecular dynamics simulations, and mutational studies (reviewed in (Goldstein and Goodey, 2021)).

We selected and filtered the DHFR sequences as explained in the methods section and generated the contact map. A comparison of the predicted contact maps using our pattern-matching approach (Fig2A) and contact maps extracted from the high-resolution structures (Fig2B) is shown in Fig. 2. Overall, the predicted contact maps follow the contacts of the true contact map. Note that for some of the domains studied (e.g., DHFR, as mentioned above), PDB structures exist in different conformations that reflect the dynamic process of their activity. These structural dynamics are also visible in the contact maps, both the one that takes the mean structure as a basis and also the one that is based on the pattern-matching approach.

**Figure 2:**
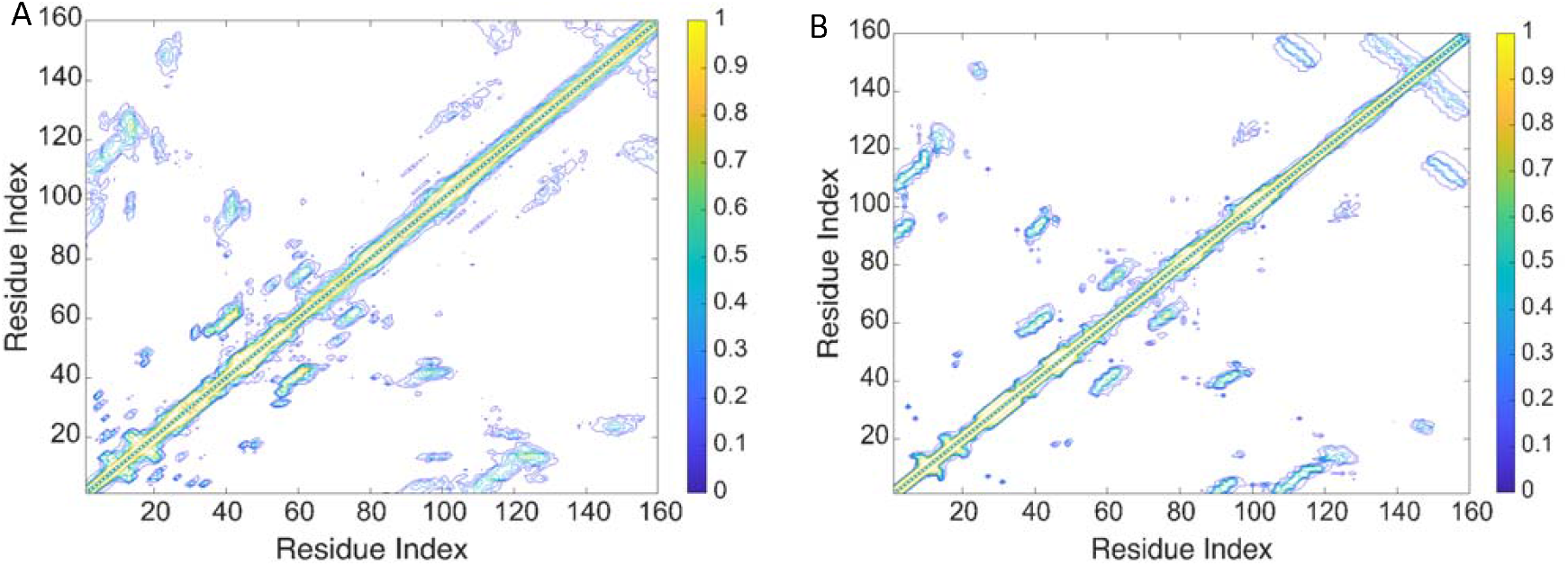
Comparison of the mean predicted and real contact maps for DHFR. (A) Mean contact maps predicted for 5895 sequences using the pattern-matching method. (B) Mean contact maps for 550 DHFR structures in the PDB.

Subsequently, we performed a dGNM analysis of the predicted contact maps. We calculated the cumulative net transfer entropy for each sequence by using Eqn. 10. For each sequence, *we identified the local maximum and local minimum values for the net transfer entropy. As an example, a plot of the cumulative net entropy transfer calculated from the predicted contact map is shown in Fig. 3 for just one DHFR sequence. Generally, negative values are labeled as entropy sinks, whereas positive values are labeled as entropy sources*.

**Figure 3:**
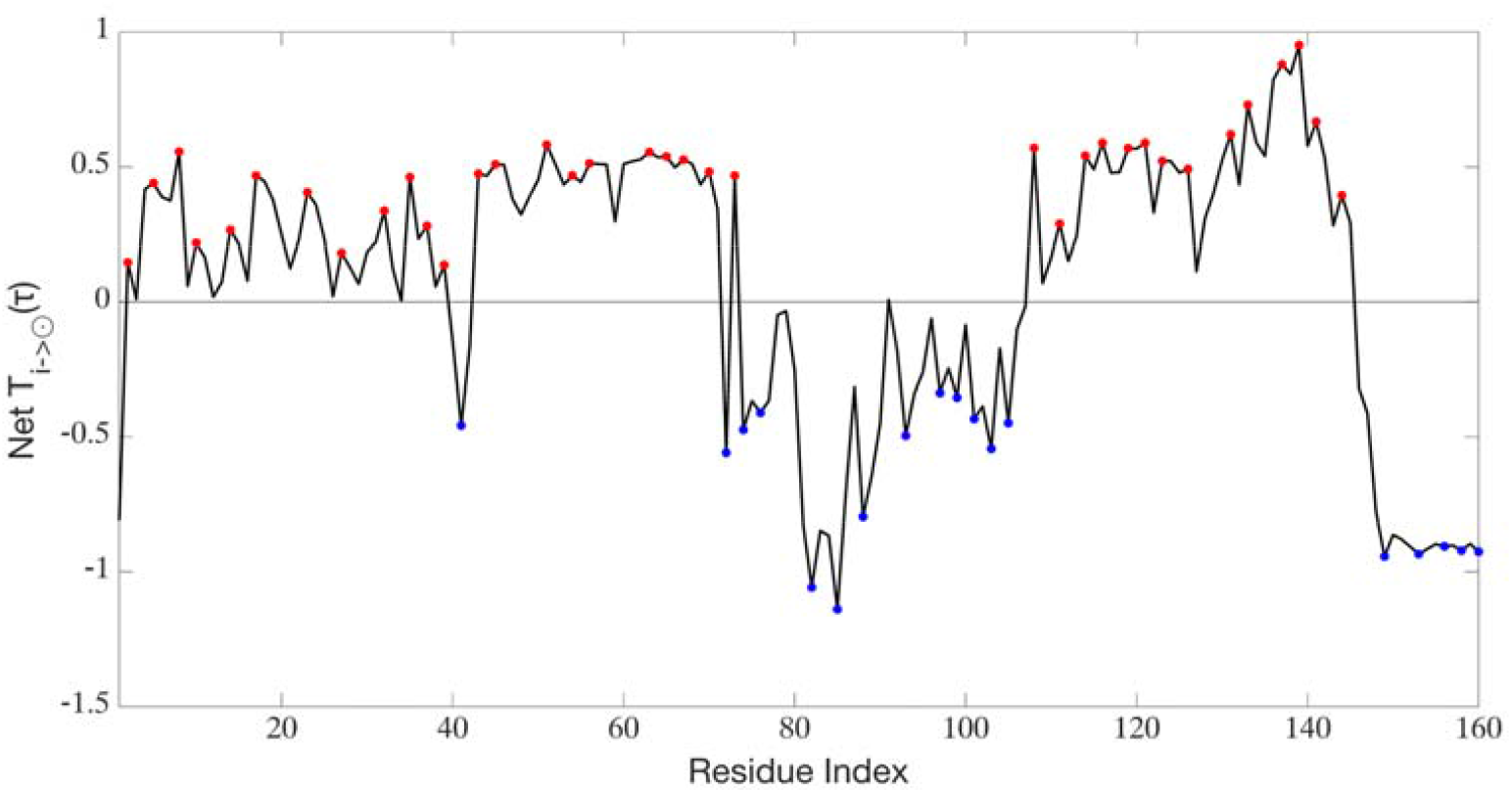
Cumulative net entropy transfer of residues calculated by using Eqn. 10 for a sample DHFR protein. Entropy source residues (local maxima) are labeled in red; entropy sink residues (local minima) are labeled in blue for the DHFR protein of the ploughfish (Gymnodraco acuticeps) (UniProt ID A0A6P8URK9).

For all DHFR sequences, we then summed up the cumulative net entropy transfer values for each source and sink residue we had detected as the local maxima and minima. We set a threshold on the total cumulative net entropy values to detect the most probable source and sink residues for the entire DHFR family. The threshold values for the total cumulative net entropy transfer for the DHFR family are indicated in Fig 3. The residues with total cumulative net entropy transfer values in the upper 15% were considered to be conserved source residues (bins labeled in red in Fig. 4) and the values in the lower 15% were considered to be conserved sink residues (bins colored in blue in Fig. 4) and thus very probably represent the residues conserved throughout the entire protein family.

**Figure 4:**
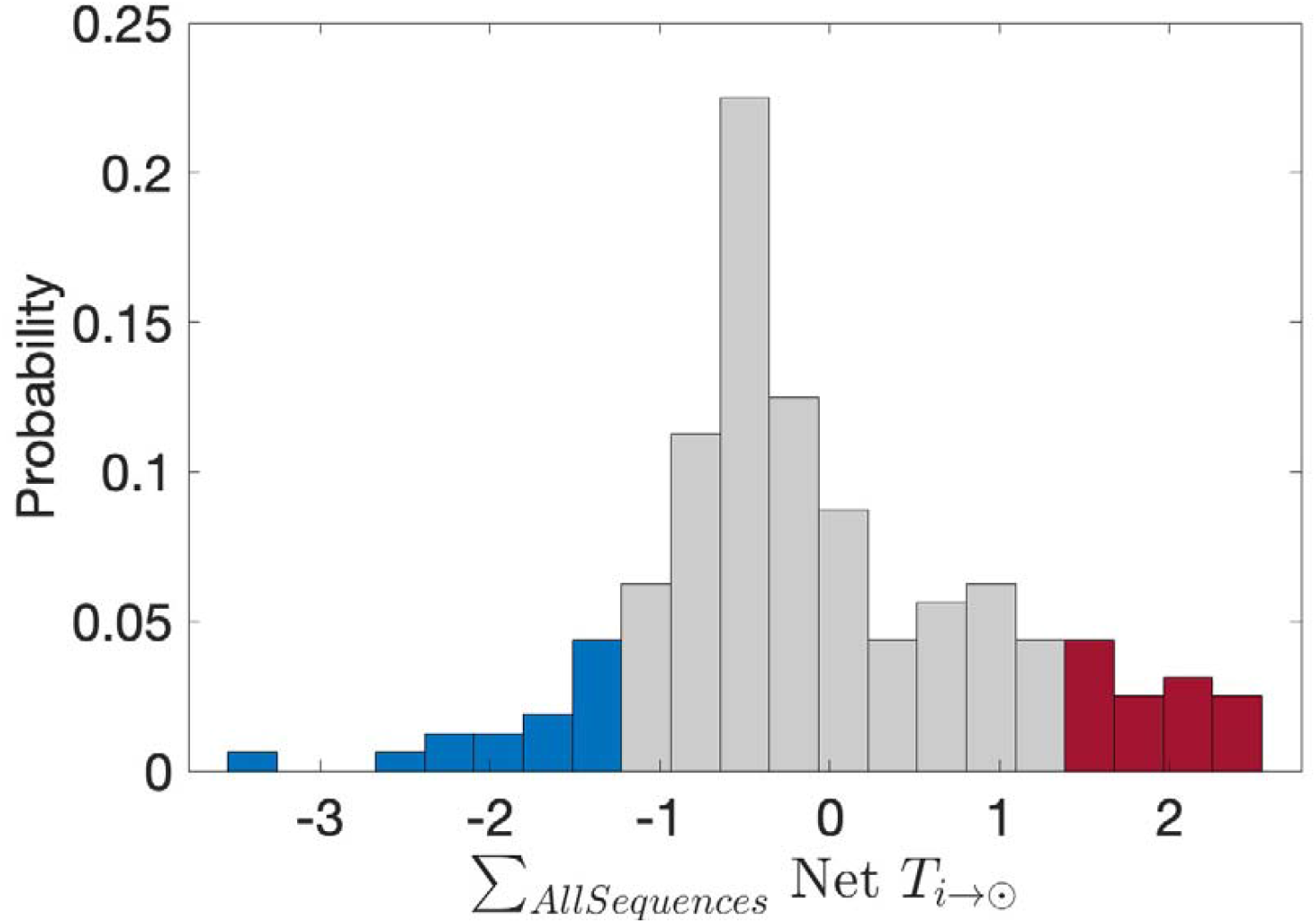
Example of a distribution plot showing the selection of thresholds for the total cumulative net entropy transfer for entropy sources and sinks for the DHFR family. The threshold for selecting the entropy sources was set at the 85 ^th^ percentile and the threshold for entropy sinks was set to the 15 ^th^ percentile.

Next, we viewed the position of the cumulative source and sink residues identified by our analysis, which revealed that loops that are critical for catalytic activity (Osborne, et al., 2001) exhibit notably high cumulative net entropy values (Fig. 5A), suggesting that a signal is transmitted from these regions to the rest of the protein. We mapped the residues with high and low cumulative net entropy values onto the structure, providing a visual representation of how this network is distributed across the domain (Fig. 5B). A comparable pattern of sink and sources was detected for the available PDB structures (Fig5. C). The majority of the information sources originate from the active site, known for its ligand-binding role. In a previous study, it was established that removing the nicotinamide adenine dinucleotide phosphate (NADP) cofactor disrupts the coordination between the FG loop and the adenosine binding domain (residues 40–50) (Van Den Bedem, et al., 2013). Our observations align with this finding, revealing that the binding of NADP induces stronger signals than DHF binding, whereby information flows from the ligand-binding sites to the Met20, FG, and GH loops, causing them to undergo conformational rearrangements and attenuation. From our analysis, we identified a network of residues that play an active role in the enzyme’s function, suggesting that the hydride transfer mechanism has been preserved throughout the course of evolution (Li, et al., 2019; Li, et al., 2021).

**Figure 5:**
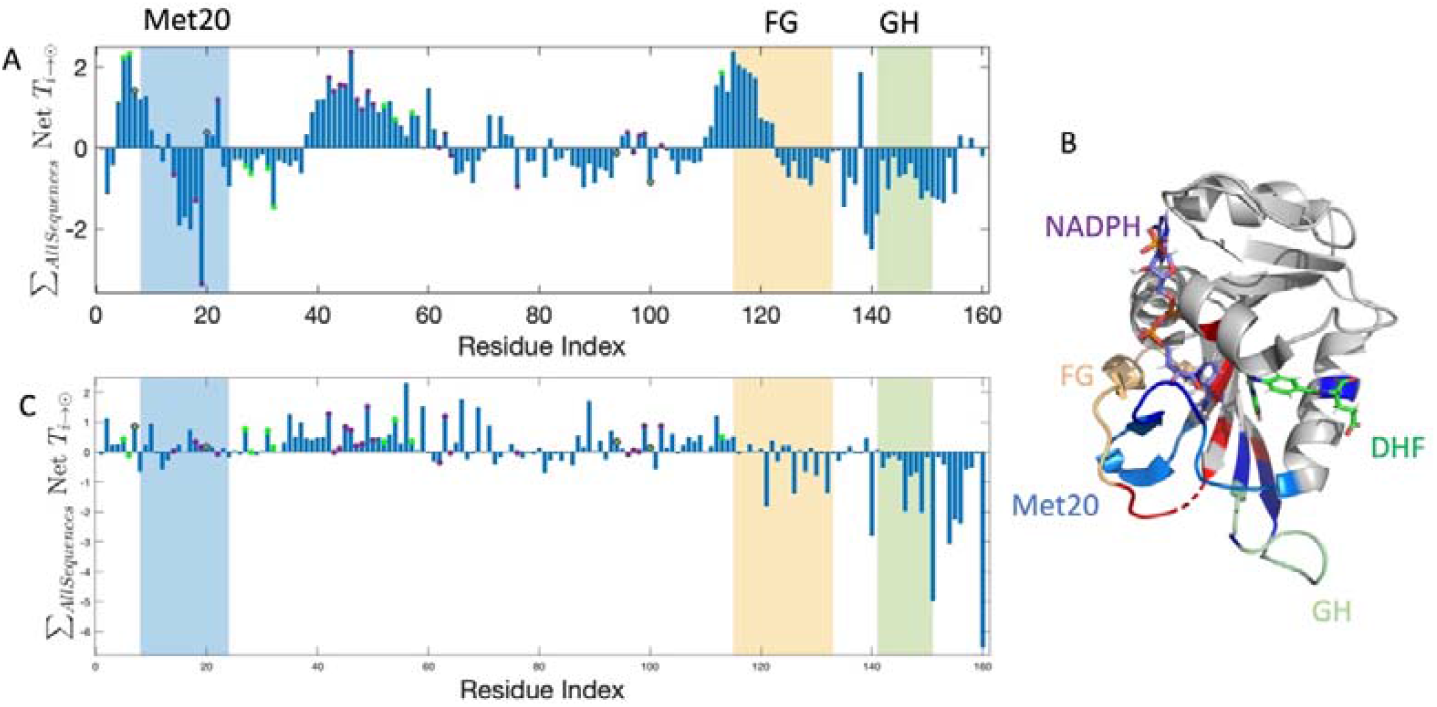
Normalized cumulative net entropy profile of DHFR. (A) Normalized summation of the cumulative net entropy values of the entropy sources (local maxima) and entropy sinks (local minima) over all selected sequences of DHFR. DHF-binding residues are labeled in green, NADP-binding residues are labeled in purple, and residues that bind both are labeled in purple-lined green dots. The regions of the Met20, FG, and GH loops are shaded on the plot with blue, yellow, and green, respectively. (B) Residues with high cumulative net entropy values (>1.72) and low cumulative net entropy values (<–1.18) are colored in red and blue, respectively, on the structure of DHFR with PDB ID 4PDJ (Wan, et al., 2014). NADP+ is colored purple and DHF is colored green. The regions of the Met20, FG, and GH loops are labeled. (C) Normalized summation of the cumulative net entropy values of the entropy sources (local maxima) and entropy sinks (local minima) for experimental DHFR structures only. Same scores as panel (A).

We also compared our results with other computational approaches; allostery pocket prediction (APOP) (Kumar, et al., 2023), MCPath (Kaya, et al., 2013) and PASSer (Tian, Jiang and Tao, 2021; Tian, et al., 2023). The two methods APOP and MCPath utilize an elastic network-based approach to detect the allosteric pockets and path residues respectively. PASSer combines pocket prediction and an automated machine learning approach to detect and label top scoring pocket residues as allosteric. The results of our method and the residues obtained by APOP and MCPath are listed in Table S1. We visualized the allosteric pockets detected by PASSer alongside the source and sink residues detected by our method by using a PyMol session (Supplementary file 1). Our results coincide with the allosteric pockets and allosteric path residues identified by the previous methods (Kaya, et al., 2013; Kumar, et al., 2023; Tian, Jiang and Tao, 2021).

Notably, nuclear magnetic resonance studies have shown that different mutations of a conserved glycine residue in the FG loop, G121, significantly slowed the transfer of hydride, except in the G121V mutant, where gating becomes kinetically significant, with three-fold faster rates of transferring hydride. Mutations of S148 in the GH loop modulate the ligand’s off-rates (McElheny, et al., 2005; Osborne, et al., 2001). Removing the central residues (residues between 16–19) of the Met-20 loop and substituting a single glycine residue result in a 500-fold reduction in the rate of transferring hydride and an increase in the rate of NADPH dissociation (Osborne, et al., 2001). X-ray studies (Sawaya and Kraut, 1997) and molecular dynamics simulations (Radkiewicz and Brooks, 2000) have suggested that the interaction of these three important loops affects how well the enzyme binds to its ligands and how fast it performs its function.

In previous studies (Altintel, et al., 2022; Skliros, et al., 2012), have shown that functionally important loops coordinate their movements through slow modes of motion. To characterize the dynamics of these loops, which we had identified as the entropy sources and sinks, even more precisely, we calculated the cross-correlations in modes 1, 3 and 10 by using the *calcCrossCorr* function of the ProDy package (Bakan, Meireles and Bahar, 2011). The cross-correlations from first three modes are shown for the sequences (Fig. 6A) and the experimental structures (Fig. 6B). The functional loops, Met20, FG and GH, are positively correlated among themselves. The cross-correlation plots for the sequences show a high level of agreement with those of the experimental structures.

**Figure 6:**
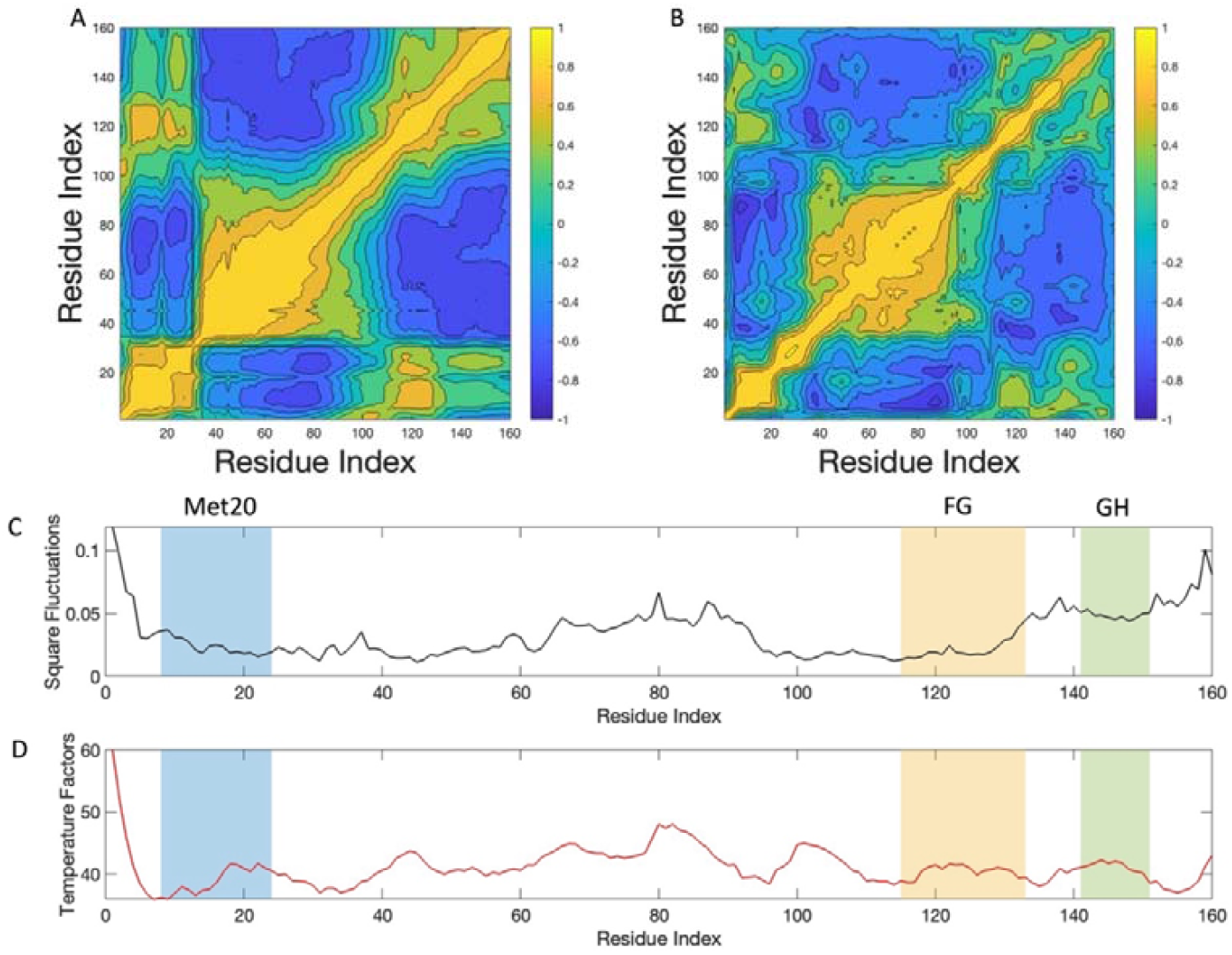
Comparison of the mean cross-correlation plots and mean square fluctuations of DHFR. (A) Mean cross-correlation plot of DHFR sequences and (B) mean cross-correlation plot of available DHFR PDB structures for the first three modes by using the calcCrossCorr function of the ProDy package. The MSE between Fig6A and Fig6B is 0.075. The cross-correlation plots for modes 1, 3 and 10 are shown in the Fig S1. (C) Mean square fluctuations of DHFR sequences and (D) mean temperature factors of available DHFR PDB structures. As in the previous Fig 5, the regions of the Met20, FG, and GH loops are shaded on the plot with blue, yellow, and green, respectively.

In Fig. 6C, the mean square fluctuations of all collected DHFR sequences, calculated by using the *calcSqFlucts* function of the ProDy package(Bakan, Meireles and Bahar, 2011), in comparison with the temperature factors of the DHFR PDB structures (Fig. 6D) are shown.

The fluctuations in Fig. 6C and the temperature factors in Fig. 6D follow a similar pattern, suggesting that our contact maps were successful in capturing the actual dynamic information of the system.

### PDZ

PDZ domains are versatile protein modules that play essential roles in mediating protein– protein interactions, particularly in the context of cellular signaling and organization. PDZ domains interact with a wide range of protein partners, including receptors, ion channels, enzymes, and scaffolding proteins. These interactions often occur within the cell membrane or at specialized cellular junctions, such as the synapse in neurons (Amacher, et al., 2020; Sheng and Sala, 2001). PDZ domains consist of approximately 90 residues that fold into a compact globular structure (Ye and Zhang, 2013). They recognize short amino acid sequences found at the C-terminus of the target proteins. Generally, PDZ domains consist of six β strands arranged in two sheets and two α helices. The peptide binds into the hydrophobic cavity between β2 and α2. PDZ domains are grouped into three types on the basis of their peptide-binding specificity: PDZ1, PDZ2, and PDZ3. PDZ2 and PDZ3 have an additional α helix at their C-terminus, which stabilizes the protein and is suggested to have a role in ligand-binding (Gautier, et al., 2018). Binding ligands into the cavity of PDZ domains leads to alterations in the motions of more distant side chains. Ligand-binding can be regulated by other proteins, for example, for the cell polarity protein Par-6. Binding of the Rho GTPase Cdc42 drastically increases the ligand-binding affinity (Whitney, et al., 2016). PDZ domains have been extensively studied for their allosteric mechanism (reviewed in (Gautier, et al., 2018) and (Stevens and He, 2022)).

To investigate PDZ domains, we first generated contact maps for the PDZ domain sequences. Note that the analysis includes only the conserved core region of all PDZ domains and does not include the third helix, which also plays a role in the allosteric regulation of the domain. It has been shown that the removal of this helix decreases the ligand-binding affinity (Chi, et al., 2012). In Fig. 7, a comparison of the predicted and real contact maps is shown. Overall, the predicted contact maps (Fig. 7A) retrieved the populated contact regions of the real contact map (Fig. 7B).

**Figure 7:**
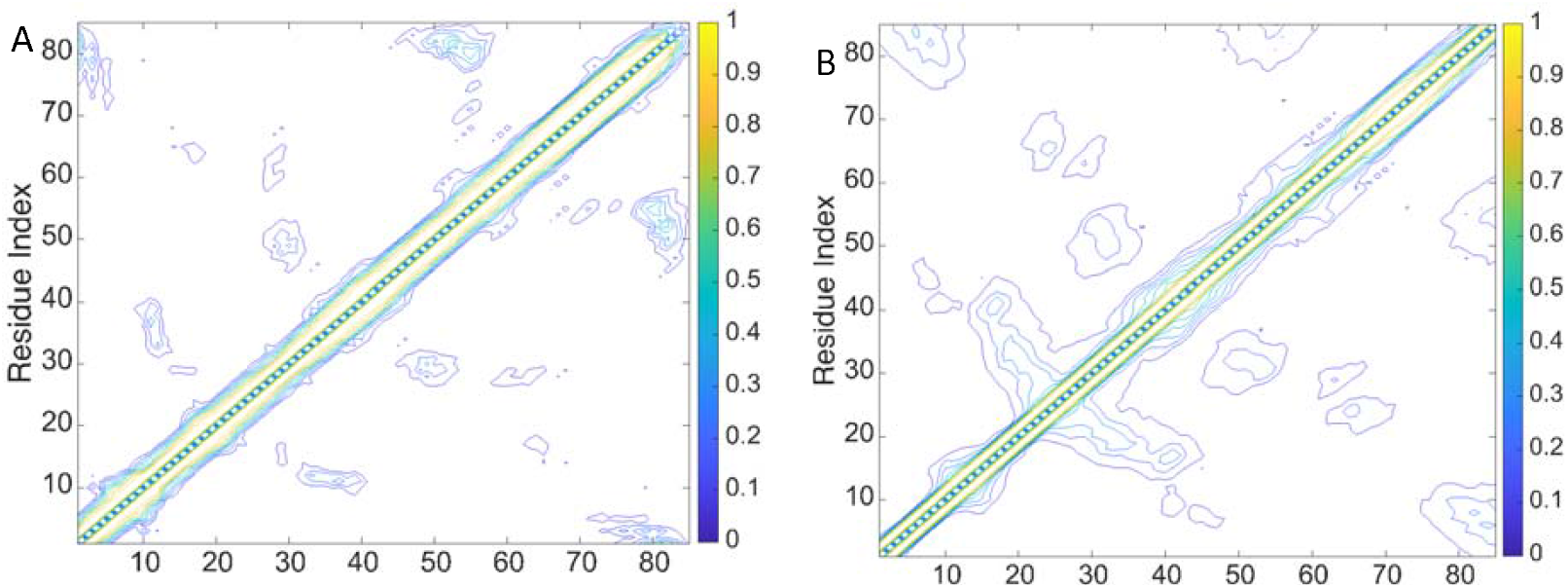
Comparison of the mean predicted and real contact maps for PDZ domains. (A) Mean contact maps predicted for 5,070 sequences using the pattern-matching method. (B) Mean contact maps for 500 PDZ structures in PDB.

As described above in detail for DHFR, we performed a dGNM analysis using the predicted contact map for the PDZ domain. We identified the conserved sources and sinks for the canonical PDZ domain through an analysis of the cumulative net transfer entropy. The sources and sinks detected (Fig. 8A) are in agreement with previous studies and with the scores calculated from the experimental structures (Fig. 8C). The majority of the signals were derived from β2, ⍰ 1 and β4, and were directed towards β1 and ⍰2, where the ligand binds the shallow pocket between β2 and ⍰2. The pattern of sources and sinks detected here matches the experimental findings (Petit, et al., 2009; Stevens and He, 2022) and likely reflects the molecular events occurring during ligand-binding in terms of allostery. This suggests that the distal residue dynamics change upon ligand-binding. In a previous study (Gerek and Ozkan, 2011), based on perturbation response scanning, the residues identified to be in the allosteric network matches with the sink and source peaks in regions of β1, β2, ⍰1, β5 and ⍰2 in Fig8A.

**Figure 8:**
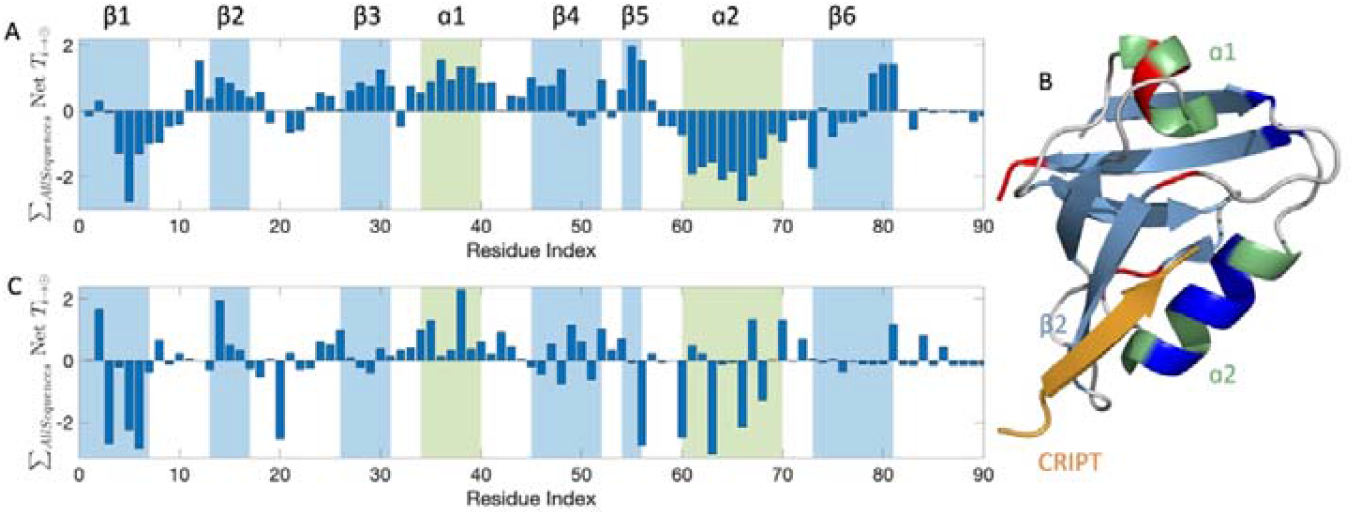
Normalized cumulative net entropy profile of PDZ. (A) Normalized summation of the cumulative net entropy of entropy sources (local maxima) and entropy sinks (local minima) over all selected sequences of PDZ. Secondary structures are colored to show β sheets in blue and ⍰ helices in green. (B) Residues with high cumulative net entropy values (>1.32) and low cumulative net entropy values (<–1.73) are colored in red and blue, respectively on the structure of PDZ3 with PDB ID 5HFF (Raman, White and Ranganathan, 2016). Note that the third ⍰ helix is not displayed. Secondary structures are colored according to Panel A. The peptide CRIPT is colored in orange. (C) Normalized summation of the cumulative net entropy values of the entropy sources (local maxima) and entropy sinks (local minima) for experimental PDZ structures only. Same scores as panel (A).

As for DHFR, we have also made a comparison with other computational approaches such as APOP, MCPath and PASSer for the PDZ domain. The results of APOP and MCPath are compared with source and sink residues detected by the current method are shown in Table S2. The allosteric path residues detected by APOP and MCPath coincides with source and sink residues detected by our method and the allosteric pockets detected by PASSer surround our source and sink residues as highlighted in the Pymol session (Supplementary file 2).

We have inspected the cross-correlation in the slow modes of motions to detect the functional regions for the PDZ domain. In Fig. S2 we present the mean cross-correlations for PDZ sequences in panels A (first mode), C (first three modes), and E (first 10 modes), and the mean cross-correlations for the experimental structures in panels B (first mode), D (first three modes), and F (first 10 modes). In the slowest modes of motions there are high positive correlations between β2-⍰1 and β5-⍰2, suggesting these regions may be functionally important. In Fig. S3A and Fig. S3B we compare the square fluctuations calculated from the PDZ sequences and the experimental structures respectively. The patterns are similar, except for fluctuations in β6. GNM models proteins as a network of connected springs based on their spatial proximity. The *N*-and *C*-terminal ends often have fewer contacts and as a consequence large amplitude fluctuations can be found among the lowest-frequency modes (Sanejouand, 2013).

### SH3

SRC homology 3 (SH3) domains are approximately 50–60 amino acids long and are involved in substrate recognition, signal transduction, cytoskeletal modification, and proliferation. They recognize proline-rich motifs with hydrophobic pockets formed by aromatic residues. Their binding site involves the region between the RT loop (between β1 and β2), characterized by conserved arginine (R) and threonine (T) amino acids, and the SRC loop (between β2 and β3). Another variable loop, the distal loop, is located on the opposite face of the domain (reviewed in (Klimov and Thirumalai, 2002; Kurochkina and Guha, 2013; Mayer and Eck, 1995; Mehrabipour, et al., 2023)). Although SH3 domains are highly conserved, relatively small, and simple, they are highly specific in their interactions. A potential approach for achieving this specificity involves adjusting the binding strength by allosteric effects, where distant residues from the binding site can influence the binding energetics for a particular substrate.

As outlined above for the other domains, we used individual sequences to generate a contact map for the SH3 domains. Overall, the populated regions of the predictions (Fig. 9A) matched the real contact map (Fig. 9B).

**Figure 9:**
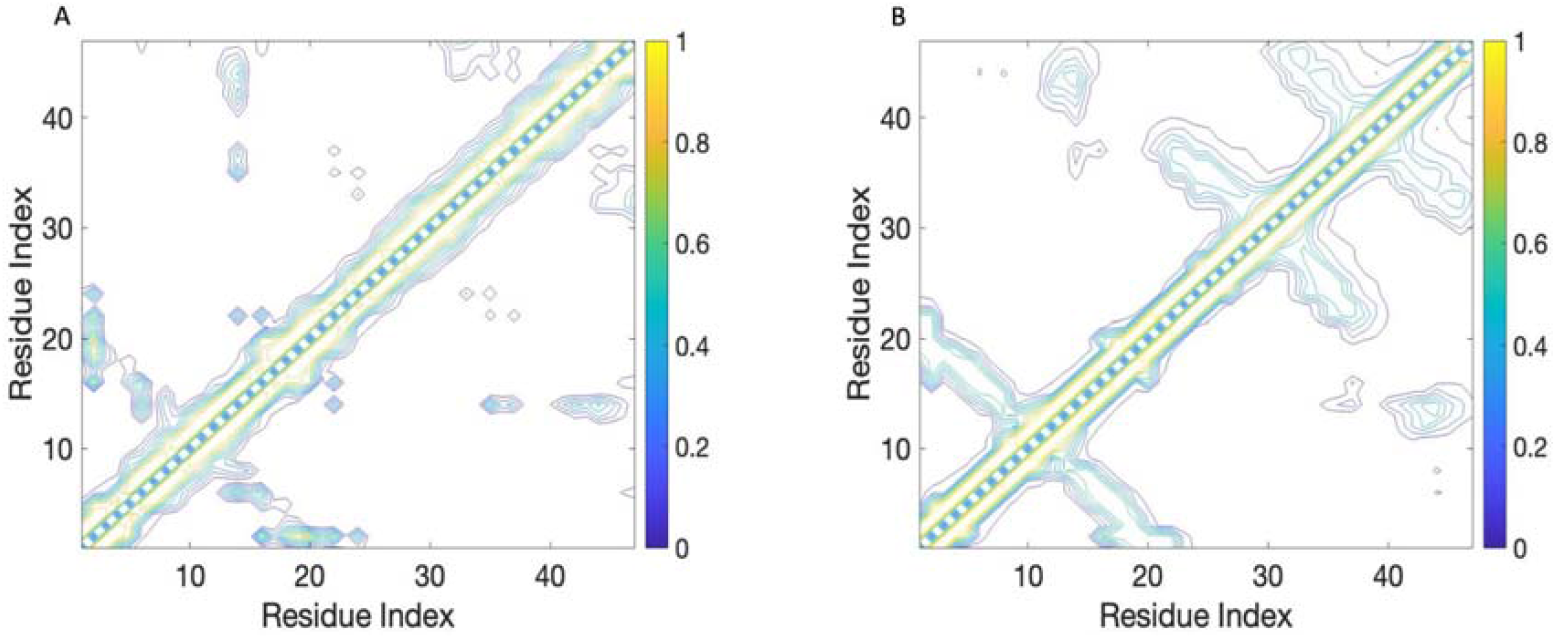
Comparison of the mean predicted and real contact maps for SH3. (A) Mean contact maps predicted for 4,674 sequences using the pattern-matching method. (B) Mean contact maps for 357 structures of SH3 in PDB. Note that some SH3 domains have slightly elongated β3 and β4 regions, which are not considered in the HMM of Pfam. Only the regions of the HMM were used for this presentation.

The normalized cumulative net entropy profile of the SH3 domains (Fig. 10A) showed that the source residues are located at the ligand-binding regions, which are distributed among the RT and SRC loops (Malagrinò, et al., 2019). Among them, two distinct residues have been proven to be the major allosteric sites for the SH3 domain (Residues 15 and 45) (Faure, et al., 2021). The distal loop, which helps stabilize the tertiary contacts, is among the regions with a strong signal source. In Fig. 10B, we demonstrate how the allosteric network’s residues are distributed through the SH3 domain and how the distant regions and the ligand-binding region connect. Our results agree with the network residues identified in previous studies (Faure, et al., 2021; Malagrinò, et al., 2019) and with the scores calculated from the experimental structures (Fig. 10C).

**Figure 10:**
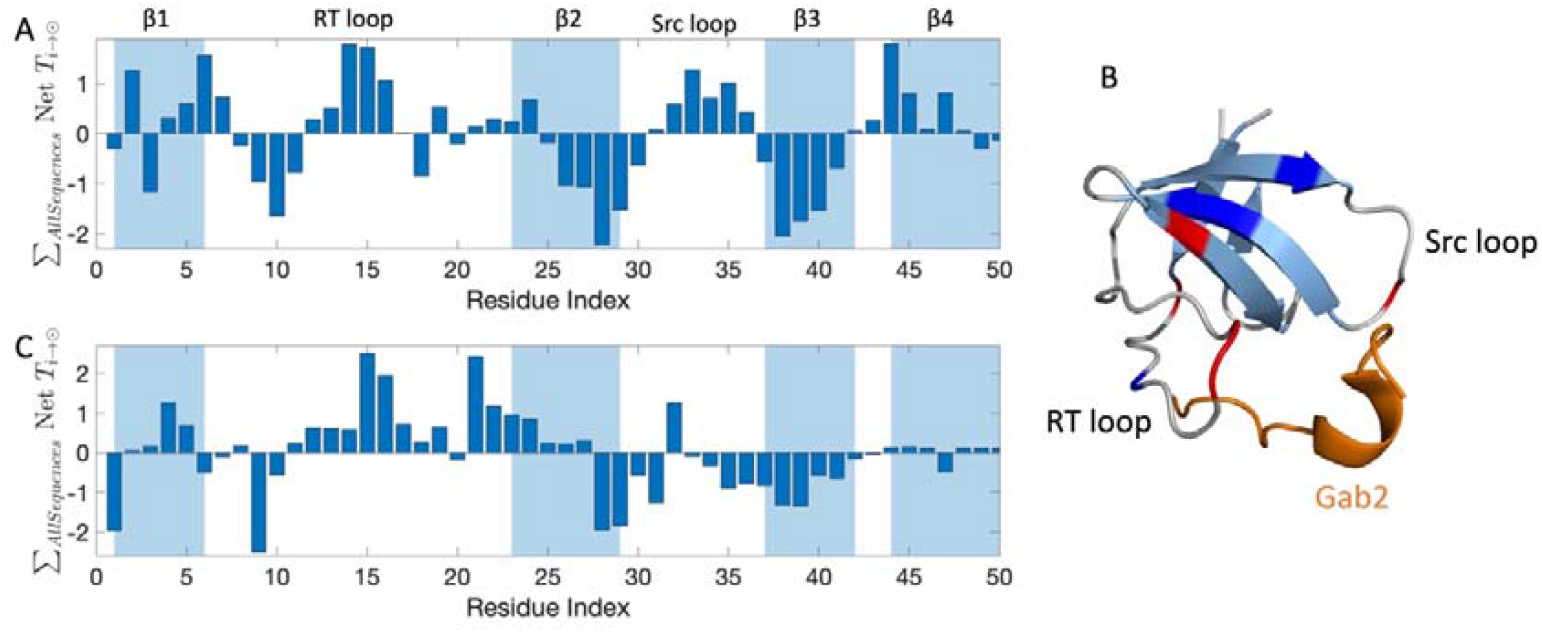
Normalized cumulative net entropy profile of SH3. (A)Normalized summation of the cumulative net entropy of entropy sources (local maxima) and entropy sinks (local minima) over all selected sequences of the SH3 domain. Secondary structures are labeled on the plot. (B) Residues with high cumulative net entropy values (>1.32) and low cumulative net entropy values (<–1.68) are colored in red and blue, respectively, on the structure of the SH3 domain with the PDB ID 2VWF (Harkiolaki, et al., 2009). The proline-rich peptide Gab2 is colored in orange. (C) Normalized summation of the cumulative net entropy values of the entropy sources (local maxima) and entropy sinks (local minima) for experimental SH3 structures only. Same scores as panel (A).

We compared our results with the other available methods (Kaya, et al., 2013; Kumar, et al., 2023; Tian, et al., 2023), as described above, for the SH3 domain. The comparison for SH3 can be found in Table S3. The comparison of the source and sink residues of our method and the allosteric pockets found by PASSer are visualized in the Pymol session in Supplementary File 3. All methods highlight the importance of the RT loop, Src loop and the N terminal of β4, as also detected by our method.

We have inspected the cross-correlation in the slow modes of motions to detect the functional regions for the PDZ domain. In Fig. S4 we present the mean cross-correlations for SH3 sequences in panels A (first mode), C (first three modes), and E (first 10 modes), and the mean cross-correlations for the experimental structures in panels B (first mode), D (first three modes), and F (first 10 modes). Mean cross-correlation plots generated from the sequences capture the high correlations of the experimental structures. In the slowest modes of motion, the RT loop is positively correlated with β1 indicating their functional importance. We also compared the square fluctuations calculated from the SH3 sequences (Fig. S5A) and the temperature factors from the experimental structures (Fig. S5B). In general, regions with higher square fluctuations correspond to higher temperature factors.

### S100

S100 proteins are a family of calcium-binding proteins approximately 90 residues long that are involved in a diverse set of functions such as enzyme regulation, cytoskeletal dynamics, signal transduction, the immune response, cell differentiation, proliferation, and migration (reviewed in (Bresnick, 2018; Fritz, et al., 2010; Marenholz, Heizmann and Fritz, 2004)). Most of them act as homodimers. Each subunit has two Ca^2+^-binding sites formed by helix– loop–helix motifs (“EF-hand type”), the C-terminal canonical EF loop is similar to the one found in other proteins such as calmodulin, and the N-terminal “pseudo-EF” is specific to S100 proteins. The two EF-hands are connected by a loop, which is critical for the interactions with the target proteins (Marenholz, Heizmann and Fritz, 2004). Normally the Ca^2+^-binding affinity of most S100 proteins is moderate (Pietzsch and Hoppmann, 2009). Interestingly, the Ca^2+^-binding affinities of both loops are different (Sivaraja, et al., 2005). Upon binding the effector proteins, the binding, the Ca^2+^-binding affinity increases through heterotropic allostery (Donato and Heizmann, 2010; Young, et al., 2022). Upon binding Ca^2+^, the domain undergoes a conformational change, exposing hydrophobic regions that facilitate interactions with other proteins (La Verde, Dominici and Astegno, 2018). The contribution of each of the two different calcium-binding loops to this cooperativity is not fully understood. Fig. 11 illustrates the underlying conformational changes caused by binding Ca^2+^.

As for the investigations above, we first established a contact map (Fig. 12C) using our pattern-matching approach and compared this with the contact maps based on high-resolution structures. As outlined above, S100 proteins undergo a larger conformational change when binding Ca^2+^ (Fig. 11). This conformational change alters the interaction pattern of the domain, as can be seen for the different contact maps of the apo (Fig. 12A) and holo states (Fig. 12B) of the protein. Remarkably, the predicted contact map (Fig. 12C) showed additional contacts between the α2 and α4 helices, and appears to represent the two conformational states, apo and holo, reflecting the dynamic nature of the allosteric regulation of S100 proteins.

**Figure 11:**
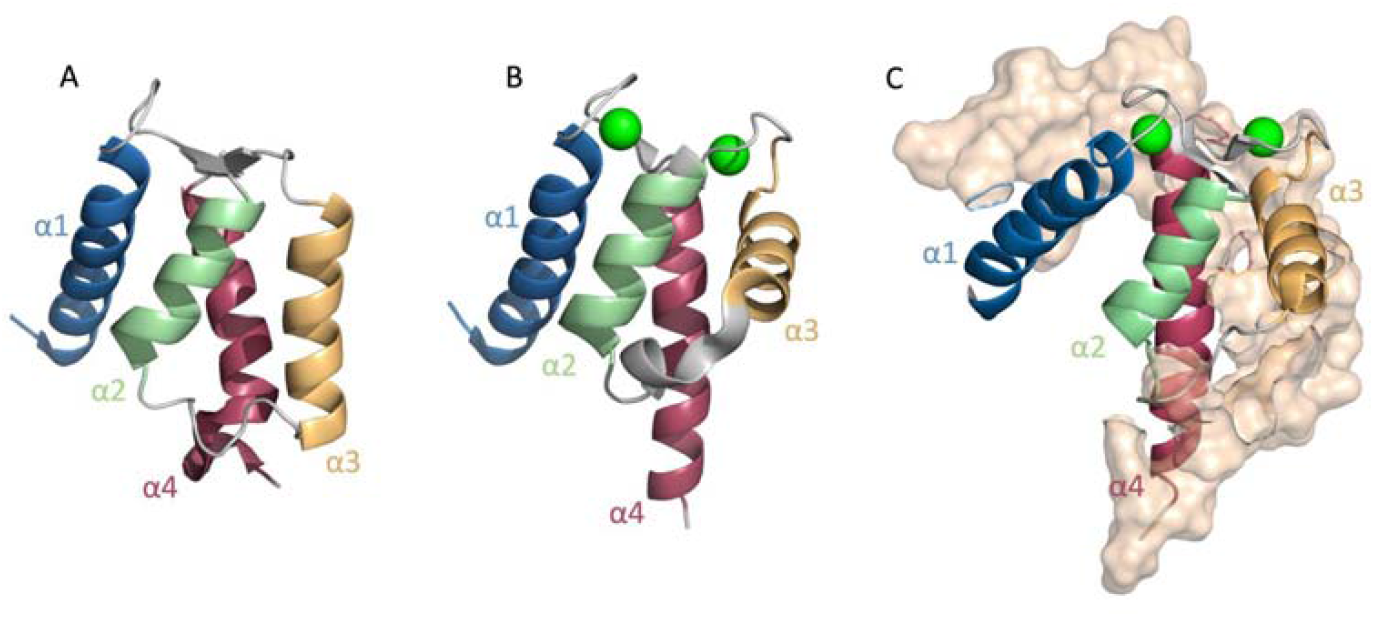
Cartoon representation of the conformational changes in the S100 domain upon binding Ca^2+^. (A) Ca^2+^ -free form (apo state) of an S100 domain (PDB ID 1K9P (Otterbein, et al., 2002)). (B) S100 domain with two Ca^2+^ ions bound (holo state, PDB ID 1K9K (Otterbein, et al., 2002)). The bound Ca^2+^ ions are shown as green spheres. Ca^2+^ -binding changes the conformation of the α3 and exposes the hydrophobic pocket for ligand-binding. (C) S100 domains with two bound Ca^2+^ ions and the possible ligand-binding regions shown as the surface, the ligands from the S100 domains of S100A4 (PDB ID 4CFR (Duelli, et al., 2014)), and S100B (PDB ID 1MWN (Inman, et al., 2002)).

**Figure 12:**
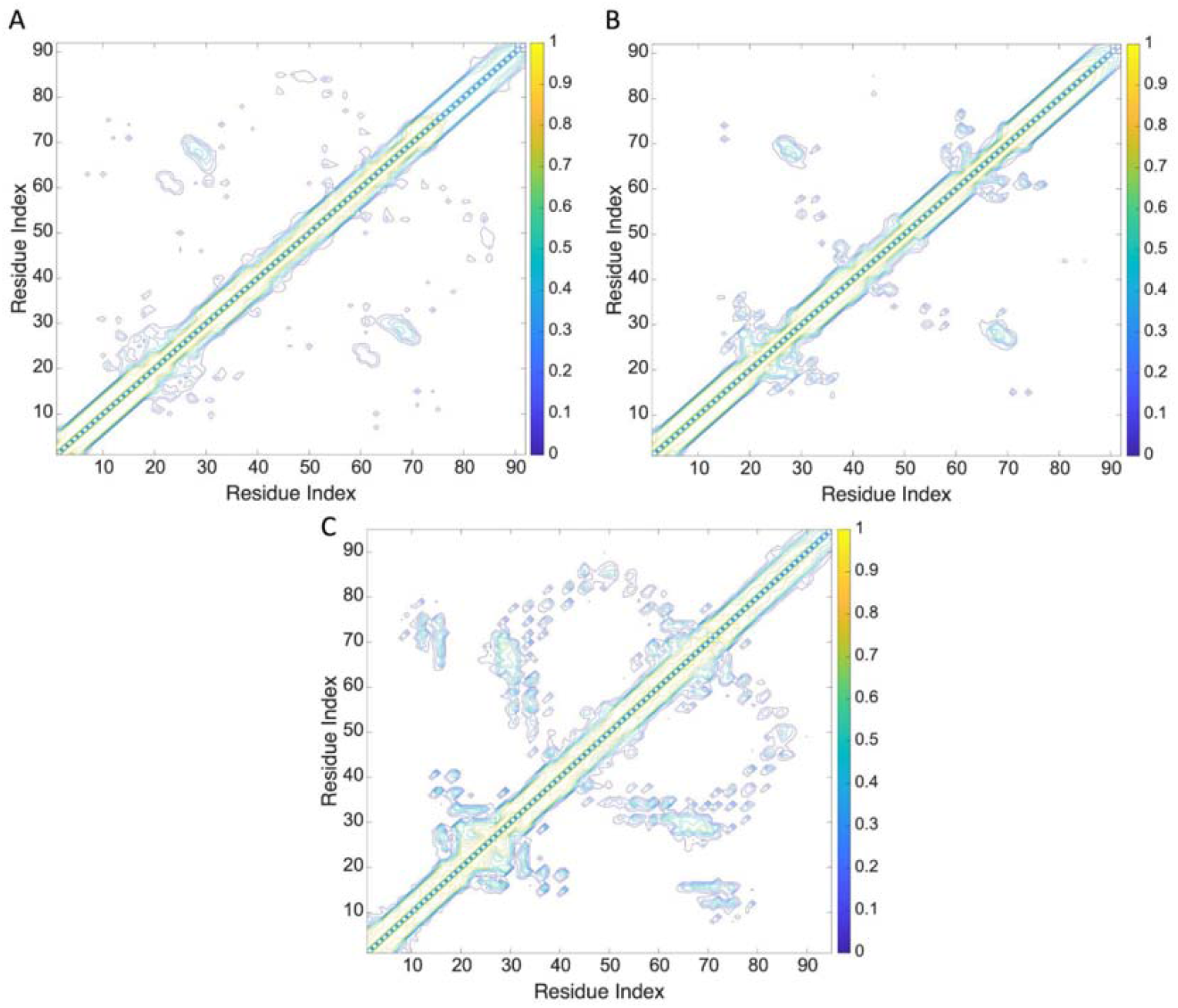
Comparison of the apo and the holo conformations of the PDB structures and the predicted contact map for S100. Mean contact maps for (A) 57 apo and (B) 108 holo structures of the S100 dimer in PDB. (C) Mean contact map predicted for 2,507 sequences using the pattern-matching method.

The normalized cumulative net entropy profile of S100 domains (Fig. 13A) showed that the binding of each Ca^2+^ induced a different type of signal. It has been discussed in previous studies that binding of Ca^2+^ to the second EF-hand is a principal trigger for the conformational switch (Wang, et al., 2024). In Fig. 13A, we can see how calcium-binding induces changes in the functionally relevant domains responsible for target recognition, such as the hinge domain between the EF-hands and the C-terminal residues. We can see how EF-I affects the overall communication in the protein after binding calcium in the EF-II.

**Figure 13:**
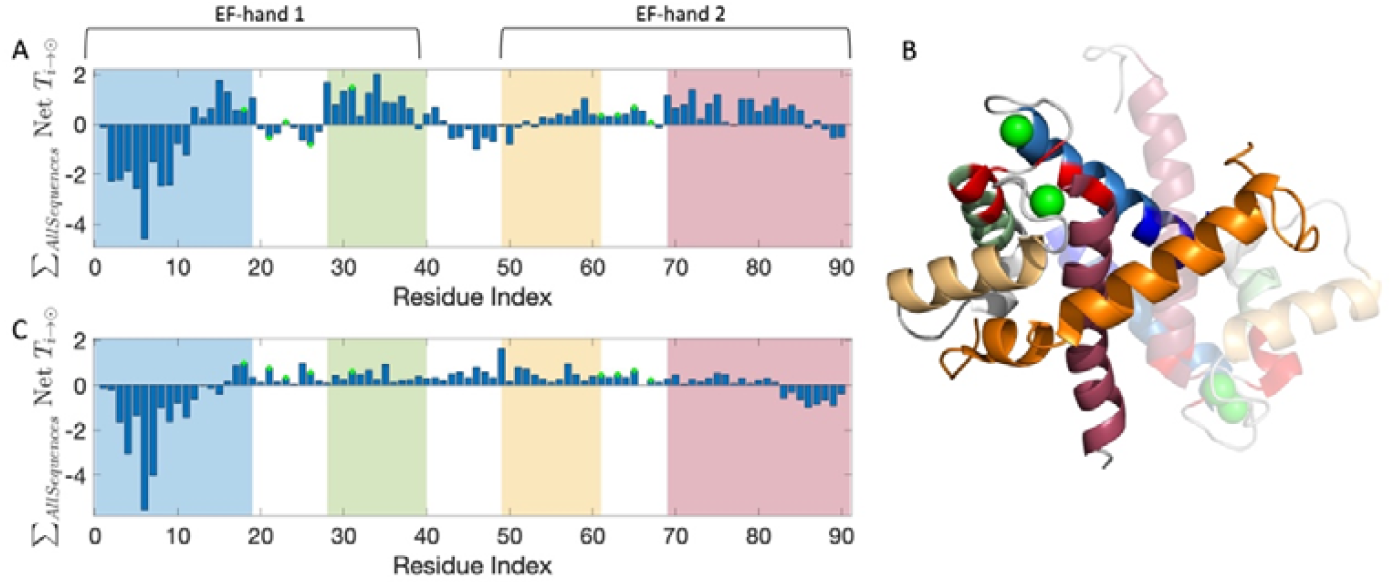
Normalized cumulative net entropy profile of S100. (A)Normalized summation of the cumulative net entropy of entropy sources (local maxima) and entropy sinks (local minima) over all selected sequences of the S100 domain. The residue ranges of the four helices are colored in order: α1, blue; α2, green; α3, yellow; and α4, red. The Ca^2+^ -binding residues in EF-hands 1 and 2 are marked with green circles. (B) Ribbon diagram of the holo structure of S100 (PDB ID 4CFR (Duelli, et al., 2014)). Residues with high cumulative net entropy values (>1.14) and low cumulative net entropy values (<–1.75) are colored in red and blue, respectively. The bound Ca^2+^ ions are shown as green spheres. The interaction partner Myosin-9 is colored orange. (C) Normalized summation of the cumulative net entropy values of the entropy sources (local maxima) and entropy sinks (local minima) for experimental S100 structures only. Same scores as panel (A).

Our findings agree with the experimental relaxation rates and molecular dynamics simulations (Wang, et al., 2024) and the profile showed in Fig. 13B. Fig. 13A captures the sink-source profiles in Fig. 13B except the behavior of few Ca^2+^ binding residues in EF-I and the loop connecting both EF hands. Since our analysis is uninformed about the conformational state of the domain, it proposes an average behavior. To elaborate more on this, we calculated the net transfer information profiles for an apo S100 domain with PDB id 1K9P and a holo S100 domain with PDB id 4CFR and the comparison is shown in Fig. S6. From Fig. S6 we can conclude that the transfer entropy profile of EF-2 changes drastically upon calcium and ligand binding.

We compared our results with the other methods (Kaya, et al., 2013; Kumar, et al., 2023; Tian, et al., 2023), as described above, for S100 domain. Table S4 shows the comparison of the source and sink residues detected by our method and the allosteric path and pocket residues detected by MCPath and APOP. The allosteric pockets detected by PASSer are compared with our source and sink residues in the Pymol session in Supplementary file 4. Overall, our method captured the residues that are found as allosteric by other methods for S100 domain.

We have also inspected the cross-correlation in the slow modes of motions to detect the functional regions for the S100 domain. The mean cross-correlations for S100 sequences are shown in Fig. S7. In the slowest modes of motion (panels C and D), the Ca^2+^ binding region of EF-I is positively correlated with the Ca^2+^ binding region of EF-II. In Fig. S8, the square fluctuations calculated from the S100 sequences (Fig. S8A) and the temperature factors from the experimental structures (Fig. S8B) are compared. Overall, the square fluctuation patterns match with the ones for the temperature factors.

## Conclusion

In this study, we present a method of exploring the conservation of allosteric mechanisms and dynamics in a diverse set of protein domains such as DHFR, PDZ, SH3, and S100. To test this method, we drew on already researched protein domains for which extensive MSAs with HMMs were available in the Pfam database and for which at least one high-resolution structure was available. We introduced pattern-matching to generate a simple 2D representation of the contact topologies of proteins directly from their sequences. This approach makes it possible to produce 2D contact maps for a large protein family without first having to create a model of the 3D structure for each sequence using Alphafold or ESMFold, which significantly minimizes the computing time required. This approach helped us analyze a large number of sequences for a given domain within a short amount of time without the need to download structures from the database or rely on GPUs on a personal computer. The contact maps created in the process largely reflect the contacts of existing 3D structures. This is a significant improvement on methods that calculate the contacts from coevolutionary correlations.

Building on Schreiber’s concept of transfer entropy(Schreiber, 2000), we identified the causal relationships and key residues in protein–protein interactions and explained the conformational changes that occur during mutation or ligand-binding and explored the dynamics of these proteins in the slow modes of motion to show their importance in terms of the function of the protein. Furthermore, our findings have important implications for the analysis of networks in newly discovered proteins, as the allosteric communication pathway and specific residues contributing to this process are preserved throughout evolution within protein domains. How well this method works for investigating other, less well-studied protein families for allosteric mechanisms needs to be shown.

This approach unveils the structural and functional intricacies of these domains, shedding light on protein–ligand interactions and allosteric communication networks, which are consistent with previous experimental findings. These insights serve to deepen our understanding of molecular mechanisms while paving the way for the development of targeted drugs and novel therapeutic interventions.

## Supporting information

Supplementary Pymol sessions

## Abbreviations

(GNM): Gaussian network model
(MD): molecular dynamics
(MSA): multiple sequence alignment
(HMM): hidden Markov models
(APOP): allostery pocket prediction
(MCPath): Monte Carlo path generation
(PASSer): Protein Allosteric Sites Server
(NADP): nicotinamide adenine dinucleotide phosphate
(MSE): Mean-squared error

## Availability of the code

The data that support the findings of this study are openly available in the Zenodo repository https://zenodo.org/records/13329929

## Acknowledgments

This work was supported by the Swiss National Science Foundation (Grant 31003A_182732 and 310030_219549 to D.F.). We thank the Division de Calcul et Soutien à la Recherche of the UNIL for access to the university’s computer infrastructure. We thank all members of the Fasshauer Laboratory and Burak Erman for helpful discussions. Special thanks to Yigit Kutlu and Turkan Haliloglu for their assistance in providing access to the MCPath servers.

## Author contributions

A.H. designed the study, performed the experiments, and analyzed the data; A.H. and D.F. wrote the paper.

