## Supplementary files for "Unveiling Conserved Allosteric Hot Spots in Protein Domains from Sequences"

Running title: Conserved Allosteric Hot Spots in Protein Domains

Aysima Hacisuleyman and Dirk Fasshauer

Department of Computational Biology

University of Lausanne

CH-1015 Lausanne, Switzerland

### Supplementary Figures

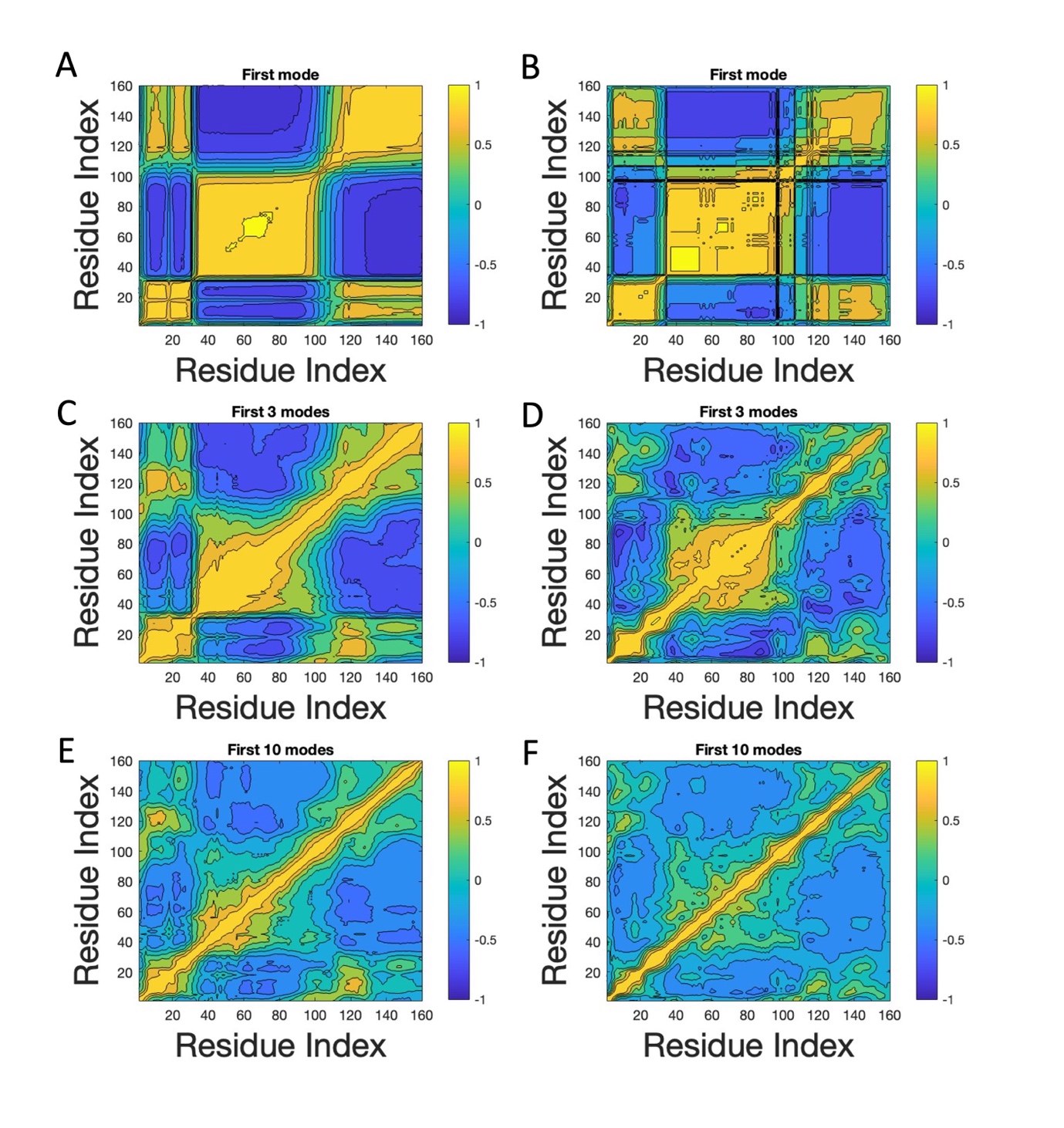

#### Fig S1: Comparison of the mean cross-correlation plots of DHFR sequences (panels A, C, E) and DHFR structures (panels B, D, F)

*Cross-correlations for panels A and B are calculated using the first modes, while panels C and D using the first three modes, and panels E and F using the first 10 modes. The mean squared error (MSE) was determined as 0.049 for panels A and B, 0.075 for panels C and D, and 0.029 for panels E and F.*

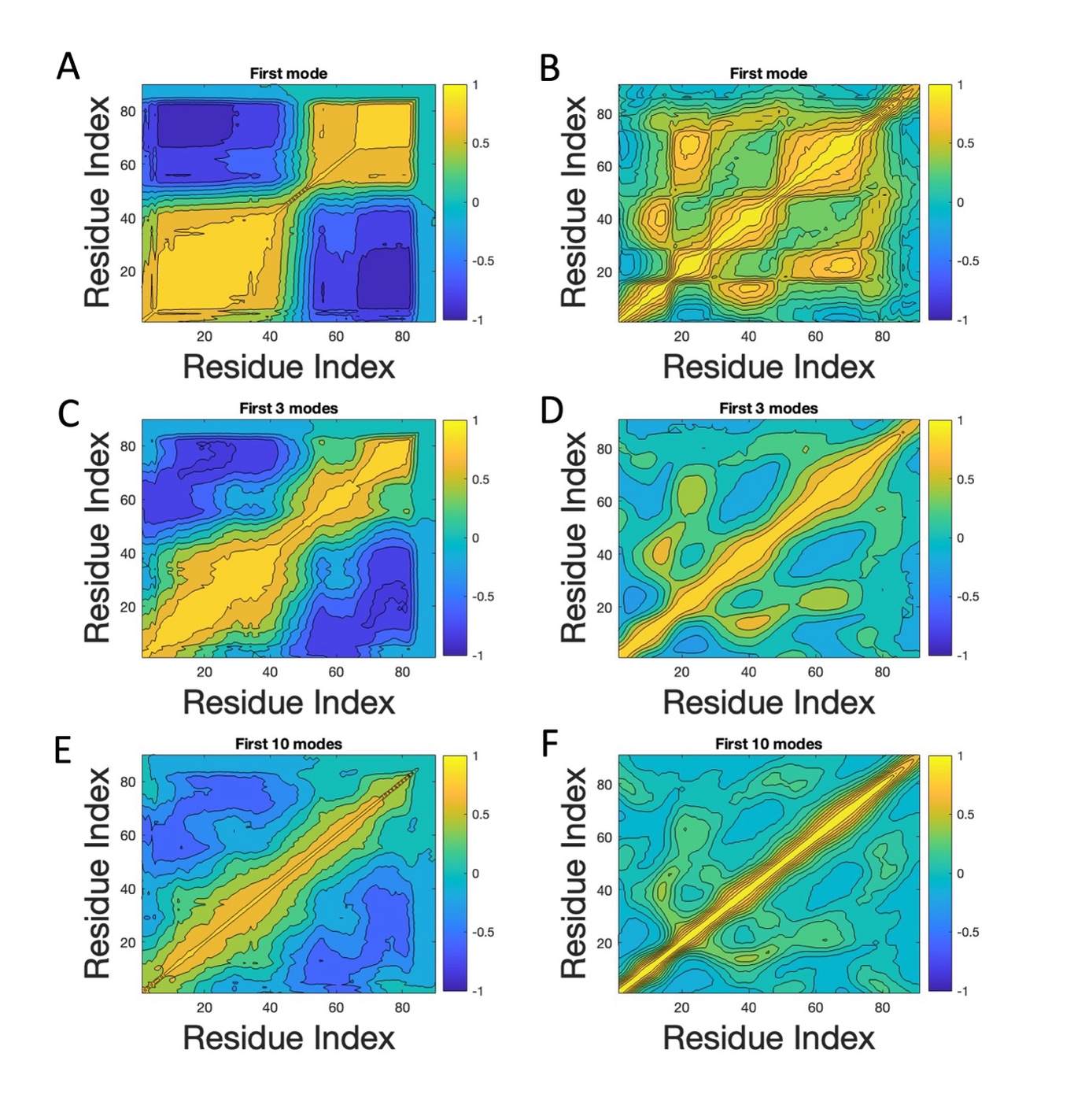

#### Figure S2: Comparison of the mean cross-correlation plots of PDZ sequences (panels A, C, E) and PDZ structures (panels B, D, F)

*Cross-correlations for panels A and B are calculated using the first modes, while panels C and D using the first three modes, and panels E and F using the first 10 modes. The mean squared error (MSE) was determined as 0.49 for panels A and B, 0.24 for panels C and D, and 0.09 for panels E and F.*

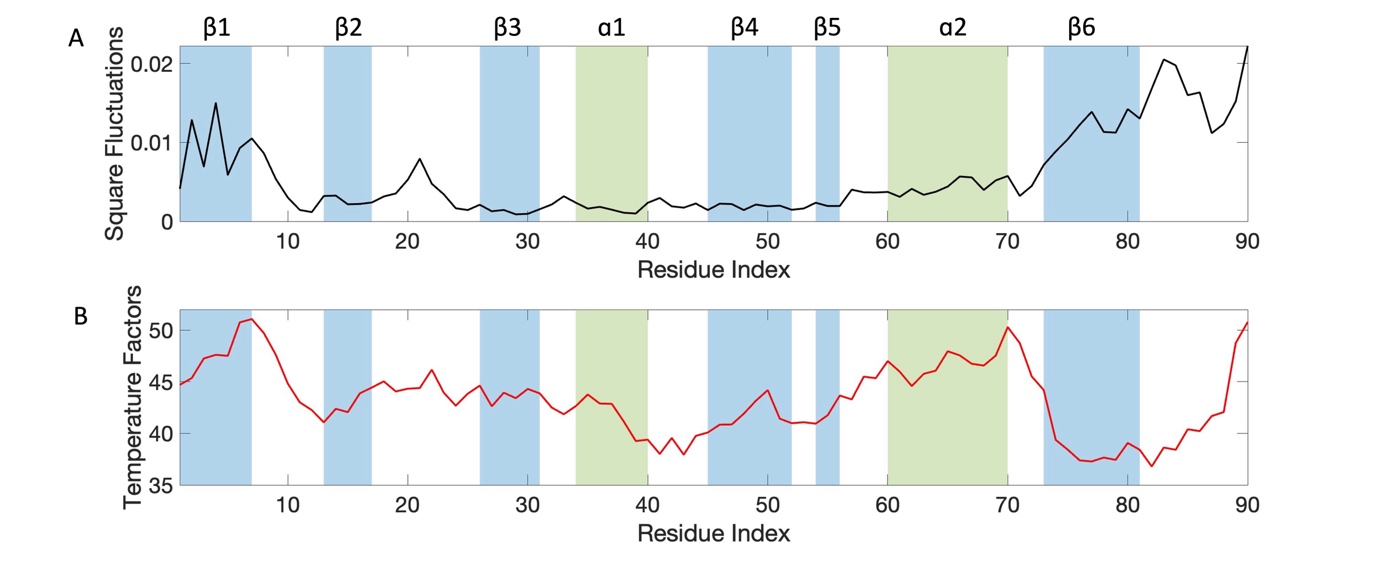

#### Figure S3: Comparison of the mean square fluctuations of PDZ.

*(A) Mean square fluctuations of PDZ sequences and (B) mean temperature factors of available PDZ PDB structures.* *As in the previous Fig. 8, the secondary structures are colored to show* β *sheets in blue and ɑ helices in green and labeled on the plot.*

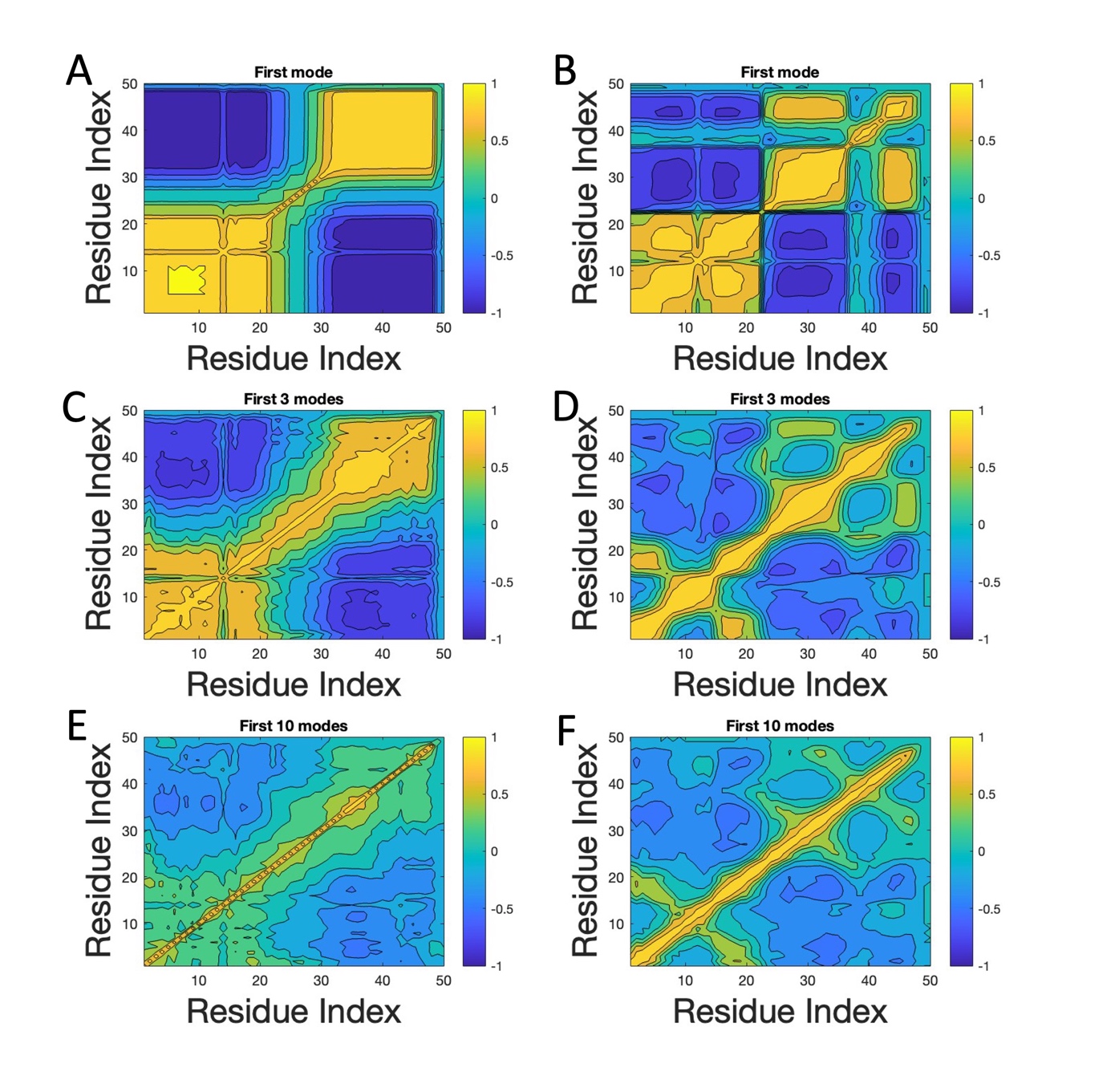

#### Figure S4: Comparison of the mean cross-correlation plots of SH3 sequences (panels A, C, E) and SH3 structures (panels B, D, F)

*Cross-correlations for panels A and B are calculated using the first modes, while panels C and D using the first three modes, and panels E and F using the first 10 modes. The mean squared error (MSE) was determined as 0.26 for panels A and B, 0.16 for panels C and D, and 0.045 for panels E and F.*

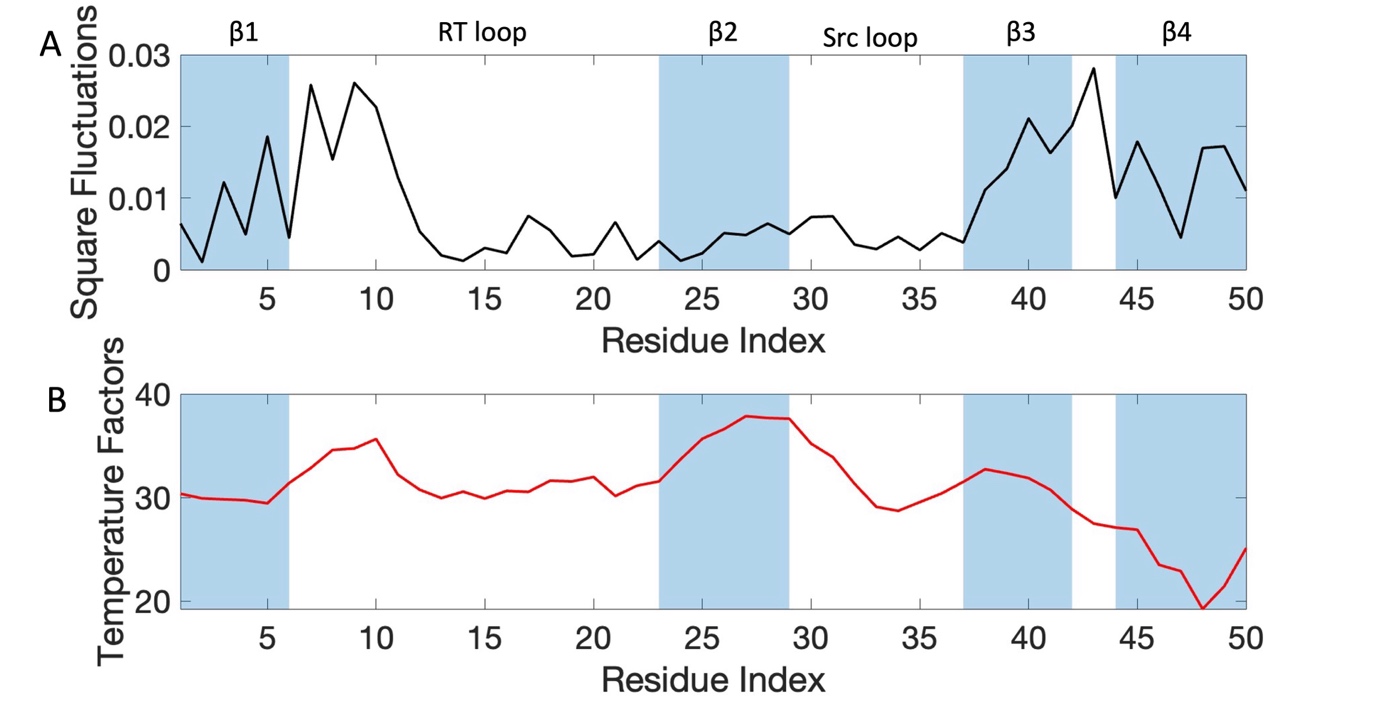

#### Figure S5: Comparison of the mean square fluctuations of SH3.

*(A) Mean square fluctuations of SH3 sequences and (B) mean temperature factors of available SH3 PDB structures. As in Fig. 10, the secondary structures are labeled on the plot.*

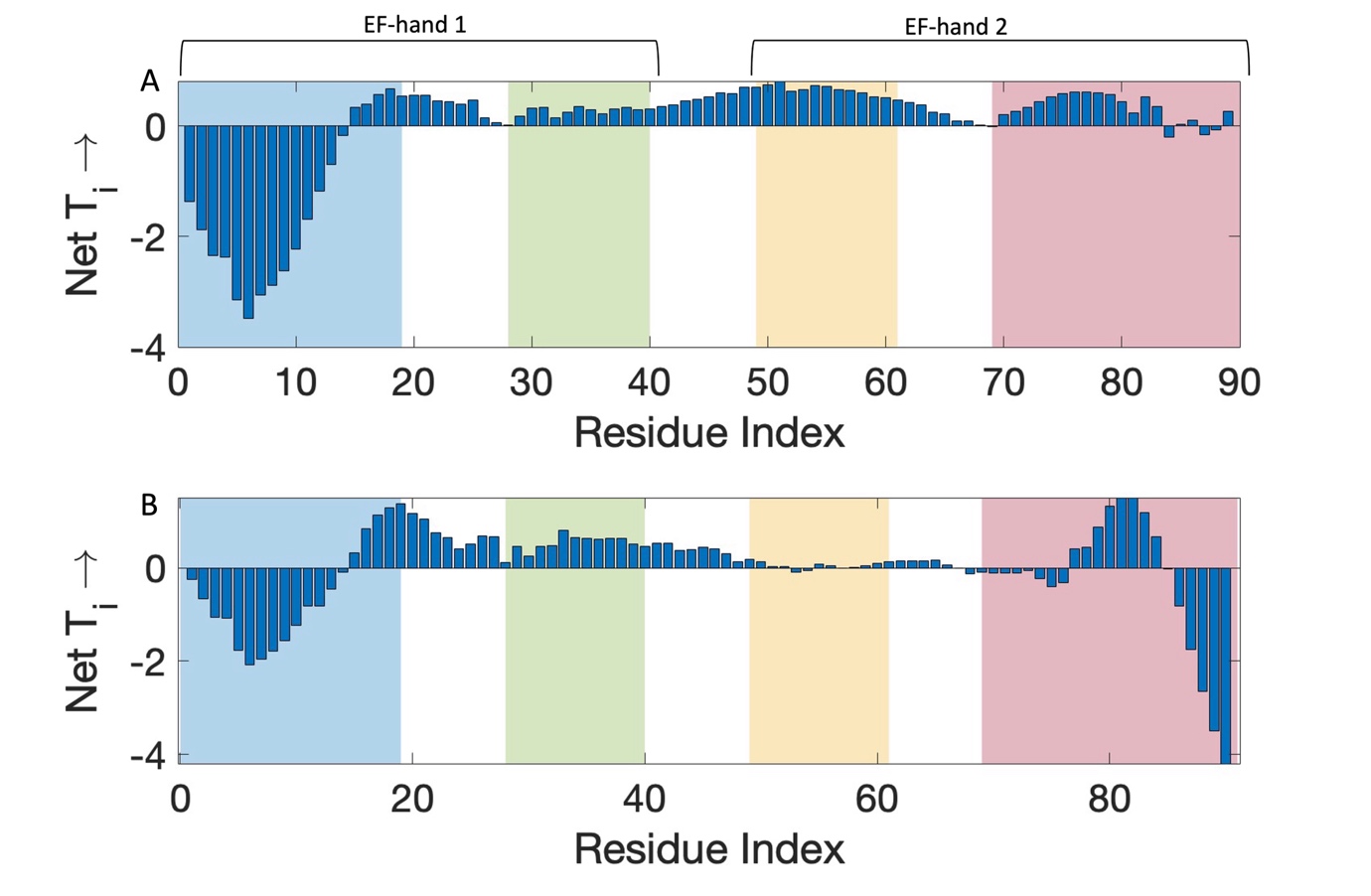

#### Figure S6: Normalized net entropy profile of apo and holo S100 domains.

*(A) Normalized net entropy of entropy sources (local maxima) and entropy sinks (local minima) of the apo S100 domain with PDB id 1KP9 (Otterbein, et al., 2002). (B) Normalized net entropy of entropy sources (local maxima) and entropy sinks (local minima) of the holo S100 domain with PDB id 4CFR (Duelli, et al., 2014). The residue ranges of the four helices are colored in order: α1, blue; α2, green; α3, yellow; and α4, red, EF hands are labeled on the plot.*

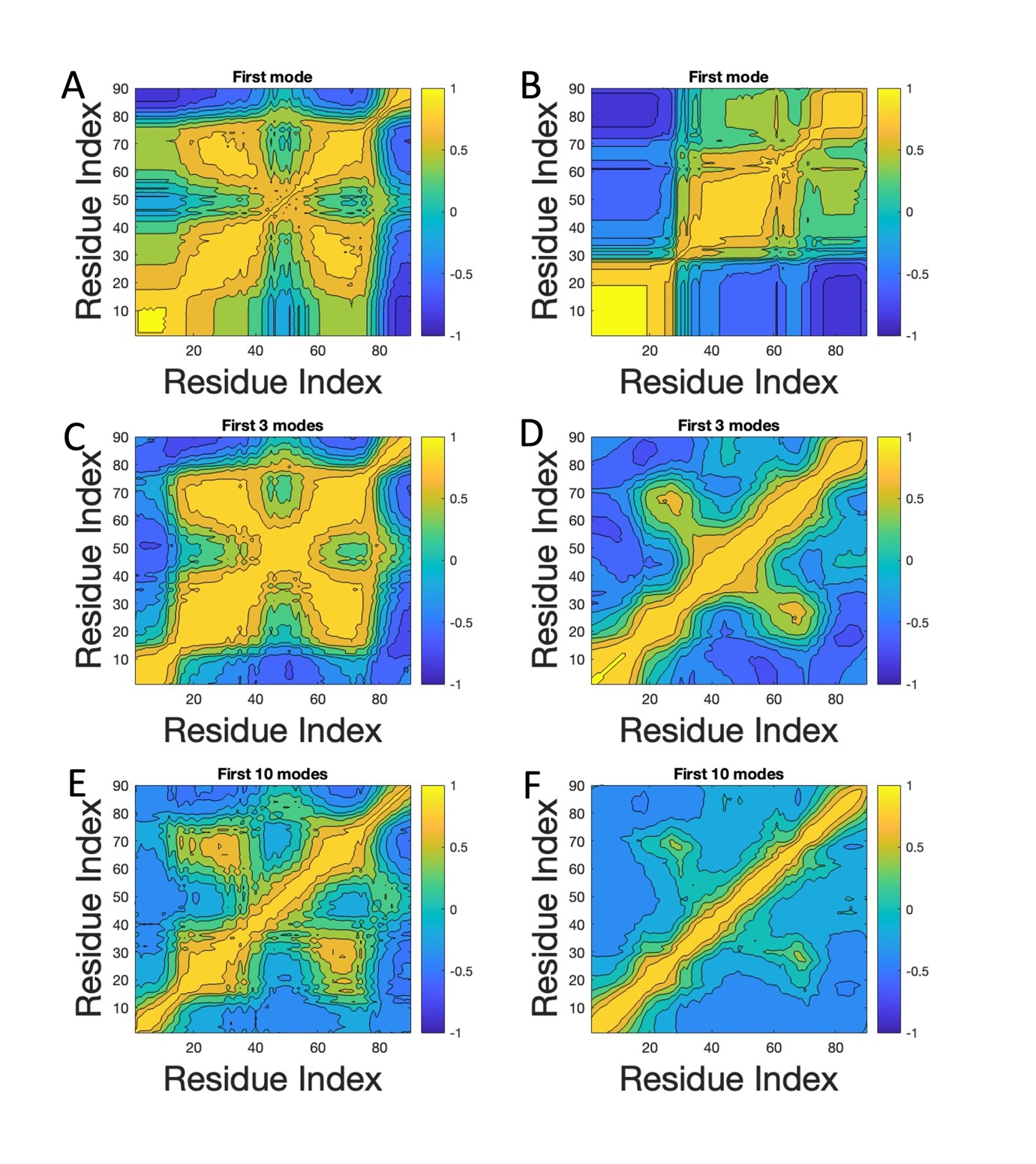

#### Figure S7: Comparison of the mean cross-correlation plots of S100 sequences (panels A, C, E) and S100 structures (panels B, D, F)

*Cross-correlations for panels A and B are calculated using the first modes, while panels C and D using the first three modes, and panels E and F using the first 10 modes. The mean squared error (MSE) was determined as 0.35 for panels A and B, 0.15 for panels C and D, and 0.07 for panels E and F.*

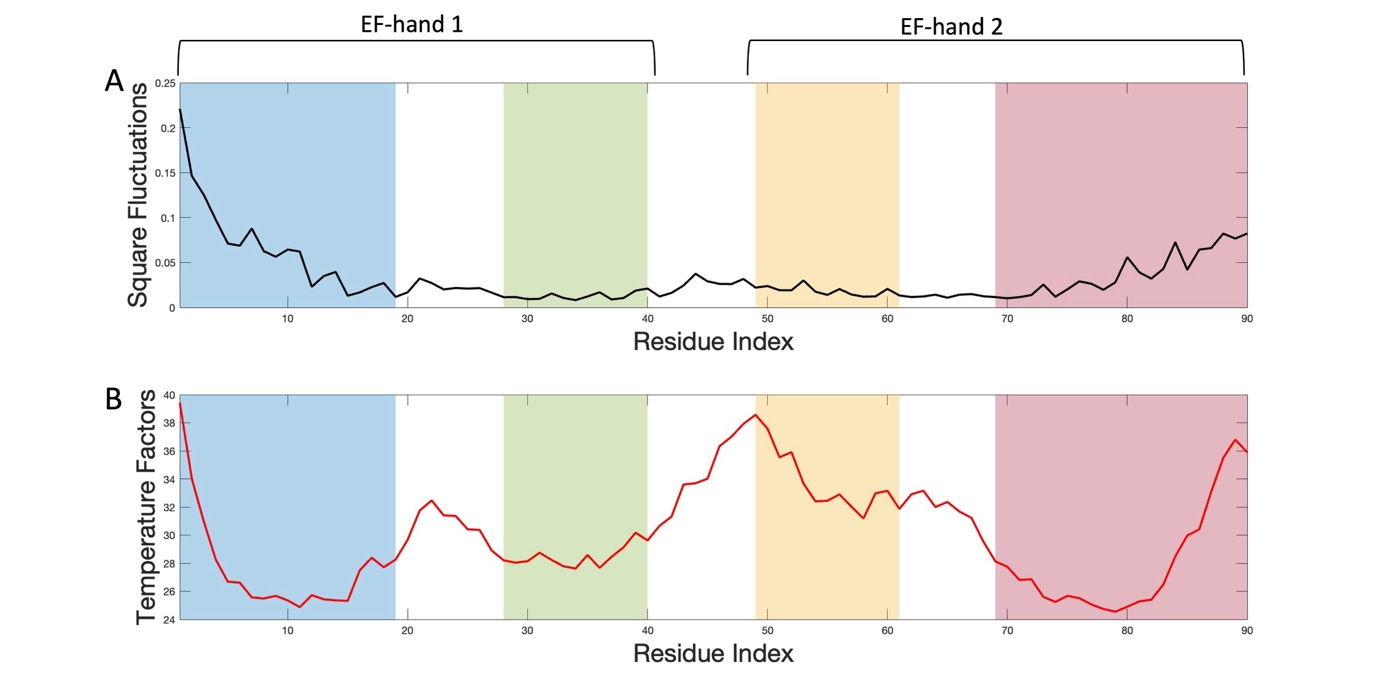

#### Figure S8: Comparison of the mean square fluctuations of S100.

*(A) Mean square fluctuations of S100 sequences and (B) mean temperature factors of available S100 PDB structures. The residue ranges of the four helices are colored in order: α1, blue; α2, green; α3, yellow; and α4, red, EF hands are labeled on the plot.*

### Supplementary Tables

***Table S1****: Comparison of source and sink residues detected by our method with Allostery Pocket Prediction (APOP) and MCPath. The structure with PDB id 4PDJ was used to generate the results with APOP, MCPath and PASSer and residue id’s are provided for 4PDJ (Wan, et al., 2014).*

| By our method | **SOURCES**  5, 6, 46, 113, 115, 116, 117, 118, 119, 138  **SINKS**  15, 16, 17, 18, 19, 32, 135, 139, 140, 141, 149, 151, 152, 153 |
| --- | --- |
| APOP | **Rank=1**  62, 64, 27, 45, 77, 57, 14, 94, 122, 100, 44, 19, 43, 30, 52, 113, 50, 99, 5, 31, 96, 6, 97, 7, 102, 123, 32, 76, 46, 124, 49, 18, 98, 20, 54, 63, 15, 28, 78, 101  **Rank=2**  63, 68, 48, 65, 74, 66, 44 |
| MCPath | **Closeness**  5, 14, 30, 34, 41, 60, 94, 100, 111, 125, 133, 153  **Betweenness**  5, 14, 18, 22, 24, 27, 30, 41, 46, 54, 57, 74, 78, 81, 94, 100, 103, 111, 125, 128, 133 153 |

***Table S2****: Comparison of source and sink residues detected by our method with Allostery Pocket Prediction (APOP) and MCPath. The structure with PDB id 5HFF was used to generate the results with APOP, MCPath and PASSer and residue id’s are provided for 5HFF (Raman, White and Ranganathan, 2016).*

| By our method | **SOURCES**  323, 347, 349, 366, 367, 391, 392  **SINKS**  316,372, 375, 376, 377, 378, 384 |
| --- | --- |
| APOP | **Rank=1**  308, 307, 351, 354, 298, 300, 301, 312, 299, 348, 341, 343, 352, 411, 314, 313, 353  **Rank=2**  380, 318, 320, 340, 384, 326, 327, 319, 383, 321, 325, 376, 323, 324, 342, 379, 322 |
| MCPath | **Closeness**  312, 325, 338, 347, 359, 367, 379, 390, 397  **Betweenness**  312, 314, 316, 318, 325, 337, 341, 354, 359, 367, 379, 392, 397, 400 |

***Table S3****: Comparison of source and sink residues detected by our method with Allostery Pocket Prediction (APOP) and MCPath. The structure with PDB id 2VWF was used to generate the results with APOP, MCPath and PASSer and residue id’s are provided for 2VWF (Harkiolaki, et al., 2009).*

| By our method | **SOURCES**  6 ,14, 15, 33, 44  **SINKS**  28, 38, 39 |
| --- | --- |
| APOP | **Rank=1**  15, 37, 38, 46, 45, 28, 44  **Rank=2**  10, 18, 41, 9, 19, 11, 20 |
| MCPath | **Closeness**  3, 9, 17, 19, 25, 36, 47, 52  **Betweenness**  3, 9, 17, 19, 25, 36, 47, 51 |

***Table S4:*** *Comparison of source and sink residues detected by our method with Allostery Pocket Prediction (APOP) and MCPath. The structure with PDB id 4CFR was used to generate the results with APOP, MCPath and PASSer and residue id’s are provided for 4CFR (Duelli, et al., 2014).*

| By our method | **SOURCES**  15, 16, 28, 30, 31, 33, 34, 72, 75  **SINKS**  3, 4, 5, 6, 7, 9, 10 |
| --- | --- |
| APOP | **Rank=1**  58, 52, 35, 45, 54, 38, 55, 51, 47, 49, 48, 39, 46, 50  **Rank=2**  9, 80, 12, 83, 79 |
| MCPath | **Closeness**  16,29, 37, 45, 58, 62, 75  **Betweenness**  12, 16, 29, 37, 42, 45, 55, 58, 72, 75, 79, 82 |

For each table, the residues which are within the upper and lower thresholds are reported for the current method. For APOP (Kumar, et al., 2023) we used a cutoff value of *7.3* Å, for MCPath (Kaya, et al., 2013) we used the *functional residue detection* option, and for PASSer (Tian, Jiang and Tao, 2021; Tian, et al., 2023) we used the *AutoML* model to generate the results.

### List of supplementary files

PASSer (Tian, Jiang and Tao, 2021; Tian, et al., 2023) results are visualized by using a Pymol Session for each domain.

DHFR; DHFR_comparison.pse (Supplementary File 1)

PDZ; PDZ_comparison.pse (Supplementary File 2)

SH3; SH3_comparison.pse (Supplementary File 3)

S100; S100_comparison.pse (Supplementary File 4)
